# Endosome-associated Rab GTPases control distinct aspects of neural circuit assembly

**DOI:** 10.1101/2025.10.09.681358

**Authors:** Katie X. Dong, Yanbo Zhang, Hui Ji, David J. Luginbuhl, Liqun Luo, Colleen N. McLaughlin

## Abstract

Neural circuit assembly relies on the precise regulation of cell-surface receptors that mediate signaling and adhesion. Endocytosis controls receptor activity and availability by internalizing and routing proteins through two main pathways: recycling back to the cell-surface or delivery to lysosomes for degradation. Rab GTPases direct receptors into these distinct pathways, but their specific contributions to circuit formation remain opaque. Using clonal analyses with null alleles, we dissected the roles of Rab-mediated trafficking to early, late, and recycling endosomes across multiple stages of circuit assembly *in vivo*. Our approach revealed that Rab5 and Rab11 regulate extensive and largely distinct developmental events, highlighting the pivotal roles of early endosome sorting and recycling pathways in circuit assembly. As neurons mature, both the spatial distribution and abundance of specific endocytic compartments change to reflect evolving trafficking demands. Our findings underscore how distinct post-endocytic trafficking fates are necessary to build neural circuits.

## INTRODUCTION

Compartmentalization is a hallmark of eukaryotic cells and is essential for nearly all aspects of cellular physiology. Organelles partition cells into spatially distinct microdomains that concentrate factors to create biochemical environments necessary for cellular function. The endolysosomal system plays a vital role in subcellular organization. This dynamic network of membrane-bound compartments integrates endocytic, biosynthetic, and degradative pathways to regulate metabolism, signaling, and protein trafficking (Settembre et al., 2013; Toshima & Toshima, 2024; Villaseñor et al., 2016; von Zastrow & Sorkin, 2021).

Neurons rely on the endolysosomal system to meet their extraordinary spatial and functional demands (Azarnia Tehran & Maritzen, 2022; Camblor-Perujo & Kononenko, 2022; Imoto & Watanabe, 2025; Yap et al., 2022). Endocytosis not only governs homeostatic processes like nutrient uptake but also underlies neuron-specific functions such as axon growth (Deshpande & Rodal, 2016; Lasiecka et al., 2014; Nishimura et al., 2003) and guidance (Pasterkamp & Burk, 2021; Zang et al., 2021), synaptic transmission (Kittler et al., 2005; Shioda et al., 2017), and neuronal polarity (Eichel et al., 2022). A major aspect of this regulation involves the selective internalization of cell surface receptors, which enables neurons to remodel their plasma membrane in response to internal and external cues. Following internalization, both soluble and membrane-anchored proteins transit to early endosomes. From there, proteins destined for degradation are trafficked through late endosomes to lysosomes (Cullen & Steinberg, 2018), while others are recycled to the plasma membrane or the Golgi apparatus (Toshima & Toshima, 2024; Grant & Donaldson, 2009). These two fates have diametrically opposed consequences: degradation causes signal termination, whereas recycling can enable signal reactivation. Thus, the endolysosomal system both shapes the neuronal surface proteome (McLaughlin et al., 2026) and plays a pivotal role in controlling signal transduction.

Rab proteins comprise a highly conserved family of small GTPases that control intracellular protein trafficking (Wandinger-Ness & Zerial, 2014; Zerial & McBride, 2001). Rabs direct the post-endocytic itineraries of internalized cargos by compartmentalizing endocytic functions and coordinating transport to distinct endocytic vesicles such as early endosomes, recycling endosomes, late endosomes, and lysosomes (Bucci et al., 1992, 2000; Rink et al., 2005; Ullrich et al., 1996; Wandinger-Ness & Zerial, 2014; Zerial & McBride, 2001). Each Rab typically localizes to a specific membrane compartment (such as Rab5 to early endosomes; Figure 1A), where it functions as a molecular switch, cycling between an inactive, cytosolic GDP-bound state and an active, membrane-associated, GTP-bound form (Stenmark et al., 1994; Zerial & McBride, 2001). In their GTP-bound state, Rabs recruit effector proteins to coordinate trafficking events such as vesicle transport, tethering, and fusion (Jordens et al., 2005; Markgraf et al., 2007; Zerial & McBride, 2001). Thus, Rabs serve as essential sorting regulators that control the abundance, localization, and signaling activities of cell surface receptors. Although endocytic Rabs have been linked to neurodevelopmental disorders (Lamers et al., 2017; Meng et al., 2023) and processes (Saxena et al., 2005; Khodosh et al., 2006; Kawauchi et al., 2010; Rodal et al., 2011; Takano et al., 2012, 2014; Falk et al., 2014; Wu et al., 2014; Taylor et al., 2015; Liu et al., 2017; Furusawa et al., 2017), much prior work has examined the function of individual Rabs in isolated developmental events or in *in vitro* systems. Thus, we lack a comprehensive understanding of how distinct Rabs contribute to the assembly of intact neural circuits *in vivo*.

**Figure 1.**
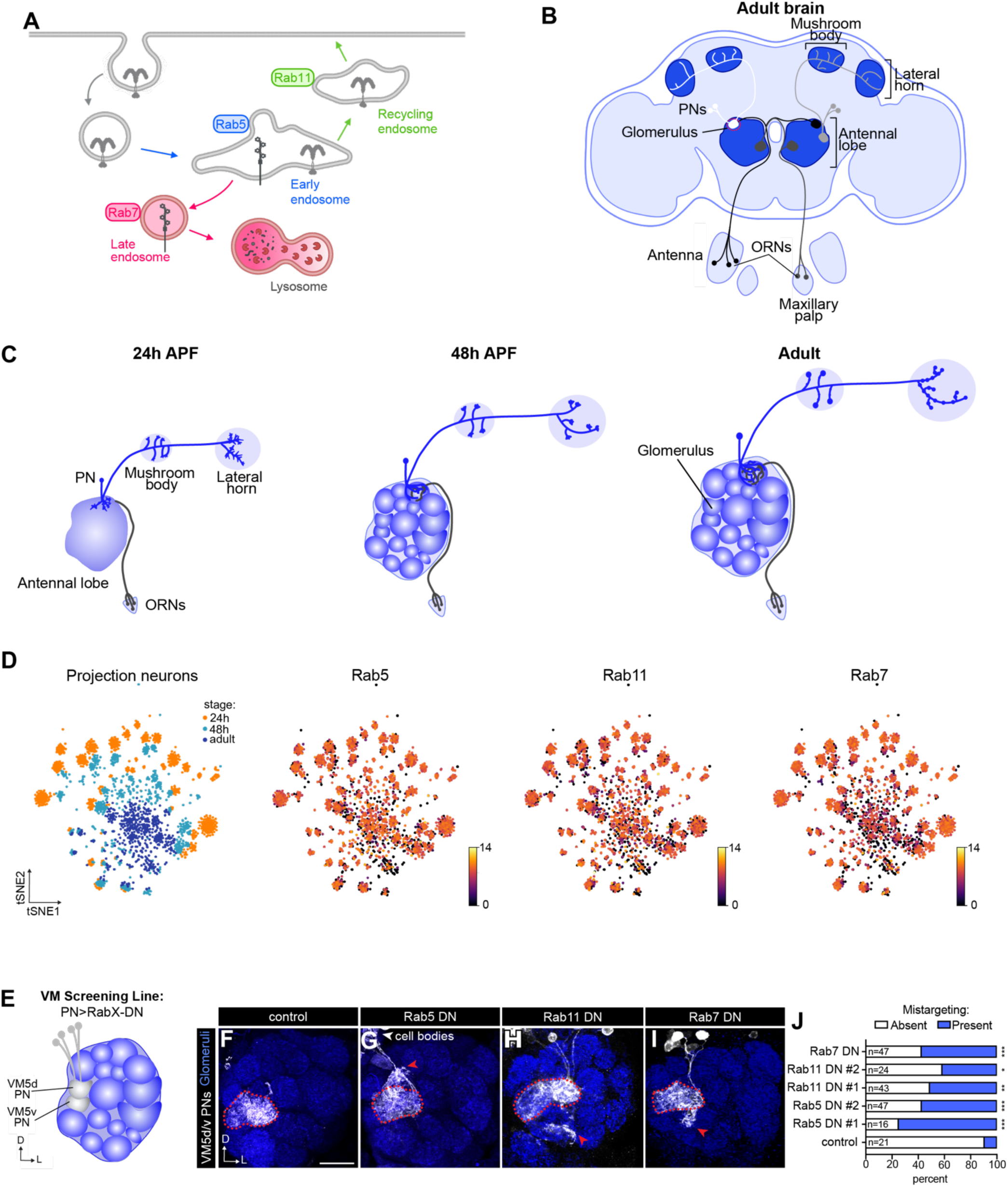
A subset of endosome-associated Rabs are expressed in olfactory projection neurons (PNs) and are involved in dendrite targeting. (A) Diagram of Rab GTPases and the endosomal compartments they predominantly localize to. (B) Schematic of the adult *Drosophila* brain, highlighting the olfactory system. ORNs, olfactory receptor neurons. PNs, projection neurons. Glomerulus on the far left depicts one-to-one matching between PNs and ORNs. (C) Schematic representing PN development at three distinct timepoints. APF, after puparium formation. (D) tSNE plots of developing PN single-cell RNA-seq (scRNA-seq) depicting stages profiled: 24h APF, 48h APF, and adult (leftmost plot) and expression of endosome-associated Rab GTPases. Expression is in log_2_(CPM +1), where CPM stands for transcript counts per million reads. scRNA-seq data are from Xie et al., 2021. (E) Schematic of ventromedial (VM) screening line used in Rab GTPase dominant negative screen. GTP-binding defective dominant negative Rabs were expressed in all PNs using *VT033006-GAL4*, and targeting of VM5d/v was monitored with *GMR86C10-LexA>LexAop-mtdTomato*. (F–I) Representative images of indicated genotypes depicting phenotypes observed in dominant negative screen. Red dotted lines outline the VM5d/v glomeruli, and red arrows denote ectopic targeting. Scale bar, 20 μm. (J) Percent of antennal lobes with mistargeting in the Rab dominant negative screen; Fisher’s exact test was used to assess statistical significance. In this and all subsequent figures: D, dorsal; L, lateral. NCad (in blue) is used to label neuropil/glomeruli. * p < 0.05; ** p < 0.01; *** p < 0.001; n.s., not significant. n, the number of the antennal lobes quantified. Images of the antennal lobe were taken of 0–5 day-old adults unless otherwise noted.

Here, we used *Drosophila* second-order olfactory projection neurons (PNs) to systematically evaluate the roles of Rab-mediated early, late, and recycling endocytic fates during circuit formation. PNs relay sensory information from the primary sensory neurons, olfactory receptor neurons (ORNs), to higher-order brain centers (Figure 1B). Assembly of the olfactory circuit begins with dendritic patterning of ∼50 distinct PN types, which are largely derived from one of two main lineages: anterodorsal (adPNs) and lateral (lPNs) neuroblasts. Within each lineage, distinct types of PNs are generated in an invariant birth order, project their dendrites to stereotyped loci within the developing antennal lobe, and their axons to higher-order brain centers (Jefferis et al., 2001, 2004; Lin et al., 2012; Marin et al., 2002; Yu et al., 2010). Subsequently, axons of each of the ∼50 ORNs enter the antennal lobe and form one-to-one synaptic connections with PN dendrites at anatomically stereotyped positions, which will eventually become glomeruli (Vosshall & Stocker, 2007; Wilson, 2013). Concurrent with ORN innervation, PN axons elaborate and refine branches and presynaptic boutons in two higher-order centers: the mushroom body and lateral horn. These developmental events unfold across a defined pupal timeline, enabling precise evaluation of dendrite targeting and axon morphogenesis within the same neuron across multiple stages (Figure 1C).

This system enabled us to compare the functions of multiple Rabs within the same cell-types spanning multiple aspects of circuit assembly. We identified endocytic Rabs critical for circuit assembly using combined transcriptomic analysis and loss-of-function screening approach. Comprehensive clonal analysis of multiple Rabs revealed that each Rab, and its related endocytic fate, is critical for distinct aspects of circuit assembly and neuronal morphogenesis. These findings demonstrate that even within a single neuronal subtype, endocytic sorting pathways are differentially employed to control separate aspects of connectivity.

## RESULTS

### Olfactory PNs express a subset of endosome-associated Rabs

The *Drosophila* genome encodes 33 Rab genes (Zhang et al., 2007), 23 of which have direct human orthologs (Chan et al., 2011; Kohrs et al., 2021), offering reduced genetic redundancy compared to the human genome, which encode 66 Rabs. Previous studies have characterized Rab GTPase expression using endogenously tagged alleles (Dunst et al., 2015; Chan et al., 2011; Jin et al., 2012; Kohrs et al., 2021). However, PN axons and dendrites reside in dense neuropil regions where they interact with the processes of other neurons and glial cells (Figure 1B, C), making it challenging to use these reagents to unambiguously define cell-type-specific expression of Rab proteins. To assess cell-type-specific expression and function of Rabs, we began by using our single-cell RNA-seq (scRNA-seq) data (Xie et al., 2021) to evaluate expression of all Rab GTPases in PNs corresponding to early (24h APF [after puparium formation]) and mid- (48h APF) development, as well as in the adult stage (Figure 1C).

We found that developing PNs express a subset of endosomal Rabs (Figure 1D, Figure 1-figure supplement 1). Most endosome-associated Rabs had one of two expression patterns. Early endosome-associated (*Rab4, Rab5*) (Gorvel et al., 1991; Kouranti et al., 2006; Simpson et al., 2004; van der Sluijs et al., 1992; Wandinger-Ness & Zerial, 2014), recycling endosome-associated (*Rab11*) (Ullrich et al., 1996; Zulkefli et al., 2019), and late endosome-associated (*Rab7*) (Rink et al., 2005) Rabs were ubiquitously and highly expressed in PNs (Figure 1D and Figure 1-figure supplement 1), whereas early-endosomal *Rab21, Rab35* (Allaire et al., 2010; Kouranti et al., 2006; Simpson et al., 2004) displayed lower overall expression (Figure 1-figure supplement 1). Finally, neither *Rab9* (late endosomes) (Lombardi et al., 1993) nor *Rab10* (recycling endosomes) (Chen et al., 2006; Etoh & Fukuda, 2019) were robustly expressed in PNs at any developmental stage (Figure 1-figure supplement 1). Our analyses indicate that PNs express a specific set of endosomal Rabs.

We also evaluated endosomal Rab expression in ORNs (McLaughlin et al., 2021), the presynaptic partners of PNs, to determine whether Rab expression patterns are a general feature of developing neurons or vary in a cell-type-specific manner. Both olfactory neuron types shared similar high expression of *Rab5, Rab4, Rab11,* and *Rab7* and low-to-absent expression of *Rab9* and *Rab10* (Figure 1—figure supplement 2). While Rab35 and Rab21 were expressed at low levels in both cell types, a smaller proportion of ORNs expressed these Rabs compared to PNs (Figure 1—supplement 2), suggesting that developing ORNs may rely less heavily on these trafficking regulators. These analyses indicate that developing olfactory neurons express a core set of endosomal Rabs.

To begin evaluating the function of these GTPases in PNs, we performed a dominant negative (DN) screen of endosome-associated Rabs. We focused on those that are highly expressed in PNs and are associated with trafficking cargos through distinct endocytic compartments. Rab2, for instance, can associate with degradative compartments but was excluded from our screen due to its primary functions in axonal transport of lysosomes (Lund et al., 2021), delivery of lysosomal proteins to the lysosomal system (Lund et al., 2018), and autophagy (Ding et al., 2019; Zhou et al., 2022). We expressed GDP-locked DN versions of each Rab, which compete with wild-type Rabs for access to guanine nucleotide exchange factors (GEFs), in the majority of PNs using *VT033006-GAL4*. To monitor dendrite targeting, we used an orthogonal driver to label VM5d/v PN dendrites within this background (shown in white; see Table S1 for complete genotype details) and N-Cadherin immunostaining to delineate glomerular boundaries (shown in blue) (Figure 1E, F).

Expressing DN versions of Rab4, Rab21, or Rab35 led to mild dendrite targeting defects typically with only a few individual dendrite branches extending beyond the VM5d/v glomerular boundary (Figure 1-figure supplement 3BA–E). Expressing DN forms of Rab5, Rab11, or Rab7, on the other hand, resulted in stronger dendrite mistargeting (Figure 1F–J) with a higher density of dendrites elaborating in incorrect glomeruli compared to Rab4, Rab21, and Rab35 defects. Although impairing each of these Rabs caused PN dendrites to target to ectopic glomeruli neighboring VM5d/v, the spatial location of mistargeting varied. For instance, disrupting Rab5 function caused dendrites to spread in dorsal, lateral, and ventral directions (Figure 1G), whereas dendrites primarily mistargeted to ventral glomeruli when Rab11 or Rab7 were disrupted (Figure 1H, I).

Rab11 is associated with slow endosomal recycling (Takahashi et al., 2012; Ullrich et al., 1996; Zulkefli et al., 2019) from recycling endosomes to the plasma membrane, whereas both Rab4 and Rab35 function in parallel fast recycling pathways directly from early endosomes to the cell surface (Allaire et al., 2010; Kouranti et al., 2006; van der Sluijs et al., 1992). The stronger phenotype observed upon Rab11 interference implies that the slow recycling route may be the predominant recycling pathway used in developing PNs. Further, among the early endosome-associated Rabs, Rab5 disruption produced stronger phenotypes than Rab21 (Figure 1G, J and Figure 1-figure supplement 3C, E). These findings are consistent with the idea that Rab5 regulates trafficking of a broad set of cargos, whereas Rab21 is thought to act more selectively on fewer cargos (Del Olmo et al., 2019; Pellinen et al., 2006; Shikanai et al., 2023). Together, these data indicate that a subset of endosome-associated Rabs is critical for PN dendrite targeting.

We wanted to leverage the mistargeting phenotypes observed upon disruption of Rab5, Rab7, and Rab11 to evaluate how distinct post-endocytic pathways contribute to circuit assembly. However, our initial analyses may have technical (e.g., late GAL4 driver expression) or biological (e.g., DN Rabs sequestering GEFs shared by multiple Rab proteins) limitations that could mask the full extent of Rab-specific contributions to this process. We circumvented this by performing clonal analyses in PNs homozygous for null alleles of *Rab5, Rab7*, or *Rab11*, which allowed us to dissect the individual roles of each Rab in PN development.

### Rab5 regulates multiple features of PN development

Early endosomes are the first major sorting station of internalized cargos. Rab5 controls early endosome biogenesis (Zeigerer et al., 2012), cargo trafficking (Bucci et al., 1992; Gorvel et al., 1991), and plays a key role in endosomal maturation toward late endosomes (Rink et al., 2005). As such, Rab5 has been implicated in dendrite arborization (Satoh et al., 2008) and targeting (Sakuma et al., 2014) in *Drosophila*, as well as in neuronal polarity (Guo et al., 2016) and dendrite development (Moya-Alvarado et al., 2018) in cultured mammalian neurons.

Given the pleiotropic function of Rab GTPases, we sought to further define neurodevelopmental roles of Rab5 by generating PN clones homozygous for a *Rab5* null allele using the mosaic analysis with repressible cell marker (MARCM) system (Lee & Luo, 1999). MARCM leverages mitotic recombination to generate homozygous mutant (or wildtype) clones marked with membrane-targeted GFP, allowing visualization of neuronal processes from individual neurons of defined genotypes against an unlabeled background (Figure 2-figure supplement 1A, B). We used this system to generate both single-PN clones for high-resolution analysis of cell-autonomous phenotypes, and larger neuroblast clones comprising multiple PNs from the same lineage, to assess multiple cell types simultaneously or potentially uncover nonautonomous effects.

**Figure 2.**
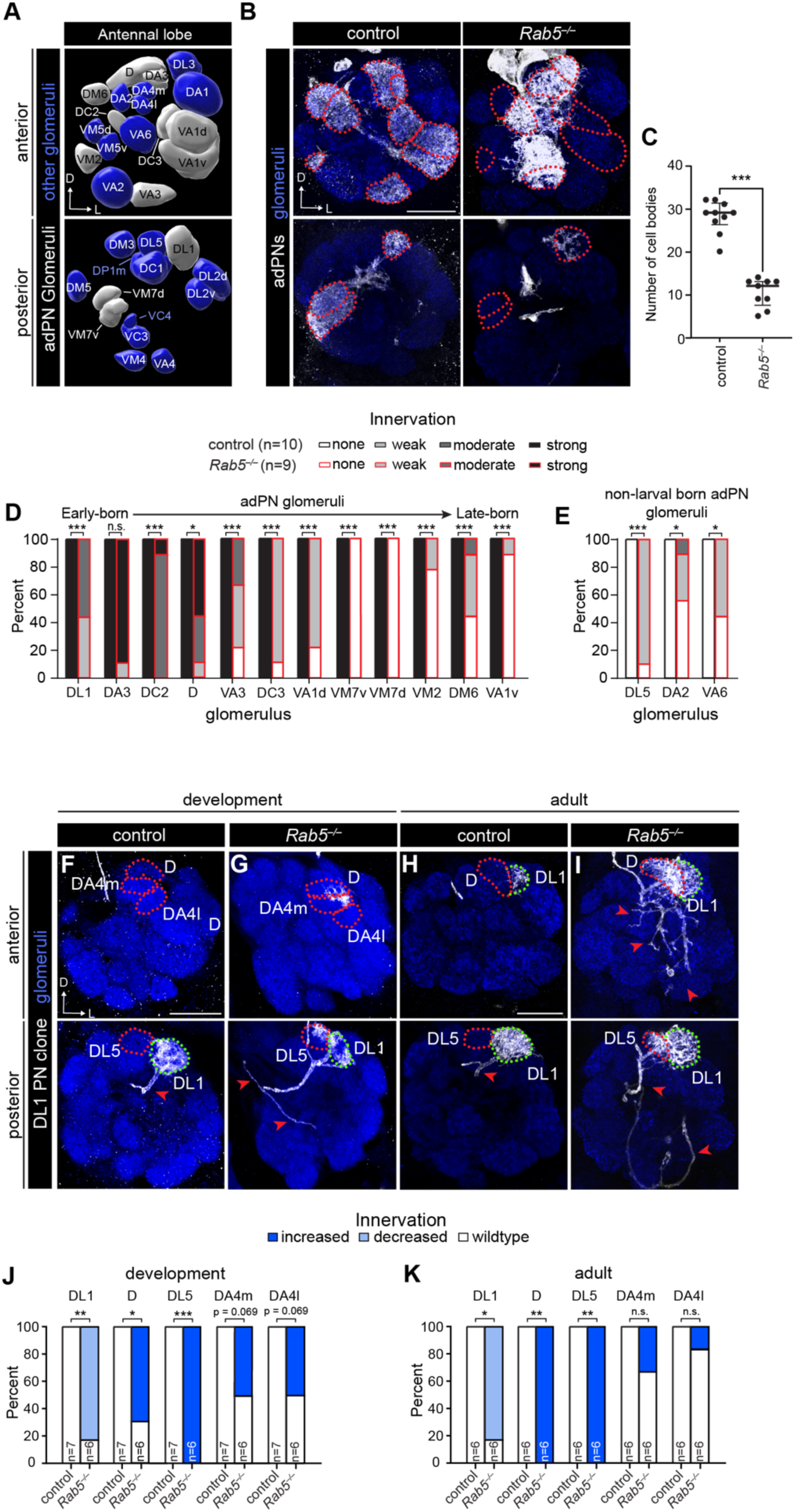
Rab5 regulates multiple features of PN dendrite development. (A) Volume rendering of a subset of glomeruli in the adult antennal lobe depicting those targeted by larval-born PNs from the anterodorsal lineage, hereafter adPNs, in grey, and other glomeruli in blue. (B) Representative images of dendrites from adPN neuroblast clones of indicated genotypes. Red dotted lines encircle adPN glomeruli. (C) Quantification of the number of cell bodies in adPN neuroblast clones in controls (n=10) and *Rab5^−/−^* mutants (n=9). Medians (IQR); Mann-Whitney U test was used to assess statistical significance. (D, E) Quantification of percent of antennal lobes with each category of dendrite innervation to adPN glomeruli (D) and glomeruli targeted by PNs other than larval-born adPNs (E). Fisher’s exact test was used to assess statistical significance. (F–I) Representative images of DL1-PN single-cell clone dendrites of indicated genotypes at 48h APF (F, G) and in the adult stage (H, I). Red dotted lines outline ectopically innervated glomeruli and green dotted lines outline DL1 glomerulus, and red arrows denote DL1-PN axon. (J, K) Quantification of percent of antennal lobes with altered dendrite innervation compared to controls in 48h APF (J) and adult (K) PNs. Fisher’s exact test was used to assess statistical significance. Scale bar, 20 µm (B, F, H).

To determine the effects of Rab5 loss on multiple PNs simultaneously, we first focused on the larval-born adPN neuroblast lineage (hereafter adPN clones) whose development and targeting patterns are well characterized (Jefferis et al., 2001, 2004) (Figure 2A). Loss of *Rab5* in adPN neuroblast clones caused a significant decrease in the number of labeled cells (Figure 2B, C), indicating that *Rab5* mutants are undergoing cell death or may have defective cell proliferation. Since PN dendrite targeting and birth order are stereotyped (Jefferis et al., 2001; Yu et al., 2010), we differentiated between these two possibilities by evaluating which glomeruli lose innervation in *Rab5* adPN clones. If PNs are dying, we would expect to observe a loss of innervation across all glomeruli, whereas proliferation defects would disproportionately affect later-born PNs. In line with the latter explanation, we observed a loss of innervation to glomeruli normally targeted by later-born adPNs compared to those that are targeted by earlier-born adPNs (Figure 2D). Beyond loss of innervation from proliferation defects, *Rab5* mutants exhibited ectopic innervation of glomeruli not normally targeted by adPN dendrites (Figure 2B, E and Figure 2-figure supplement 1C). Since only adPNs were labeled and manipulated in this analysis, these data indicate that *Rab5* also regulates dendrite targeting specificity. Taken together, Rab5 is critical for both neuroblast proliferation and dendrite targeting.

We next tested the cell-autonomous role of Rab5 in PN dendrite targeting. By generating DL1-PN single-cell clones and quantifying dendrite targeting at a stage when glomerular boundaries have just formed during development (48h APF), we could directly observe targeting errors close to when they arise. Loss of *Rab5* in DL1-PN clones resulted in ectopic innervation into the neighboring glomeruli D and DL5 as well as reduced innervation to the DL1 glomerulus (Figure 2F, G, J), consistent with a previous study (Sakuma et al., 2014). Dendrite mistargeting persisted into adulthood (Figure 2H, I, K), indicating that these defects are not corrected by removal of aberrant projections. Although the deficits did not reach statistical significance, we also observed consistent mistargeting to DA4m and DA4l glomeruli (Figure 2G, I, J, K) in Rab5 single-cell clones. Thus, Rab5 cell-autonomously regulates PN dendrite targeting.

### Rab5 has target-specific effects on axon development

PN axons exit the antennal lobe and project to the mushroom body and lateral horn (Figure 3A) where they elaborate branches and form presynaptic boutons. Rab5 has been implicated in regulating axon elongation (Falk et al., 2014; Sakuma et al., 2014), fasciculation (Wu et al., 2014) and guidance (Sakuma et al., 2014; Wu et al., 2014); however, its roles in other developmental processes are largely undefined. Our ability to generate DL1-PN mutant clones enabled us to evaluate the function of Rab5 in axon growth and guidance as well as finer-scale axon morphology. Consistent with previous work (Sakuma et al., 2014), 40% of the DL1-PN clones terminated in the vicinity of the antennal lobe rather than projecting to higher-order regions (Figure 3B, right), indicating that Rab5 is important for axon guidance.

**Figure 3.**
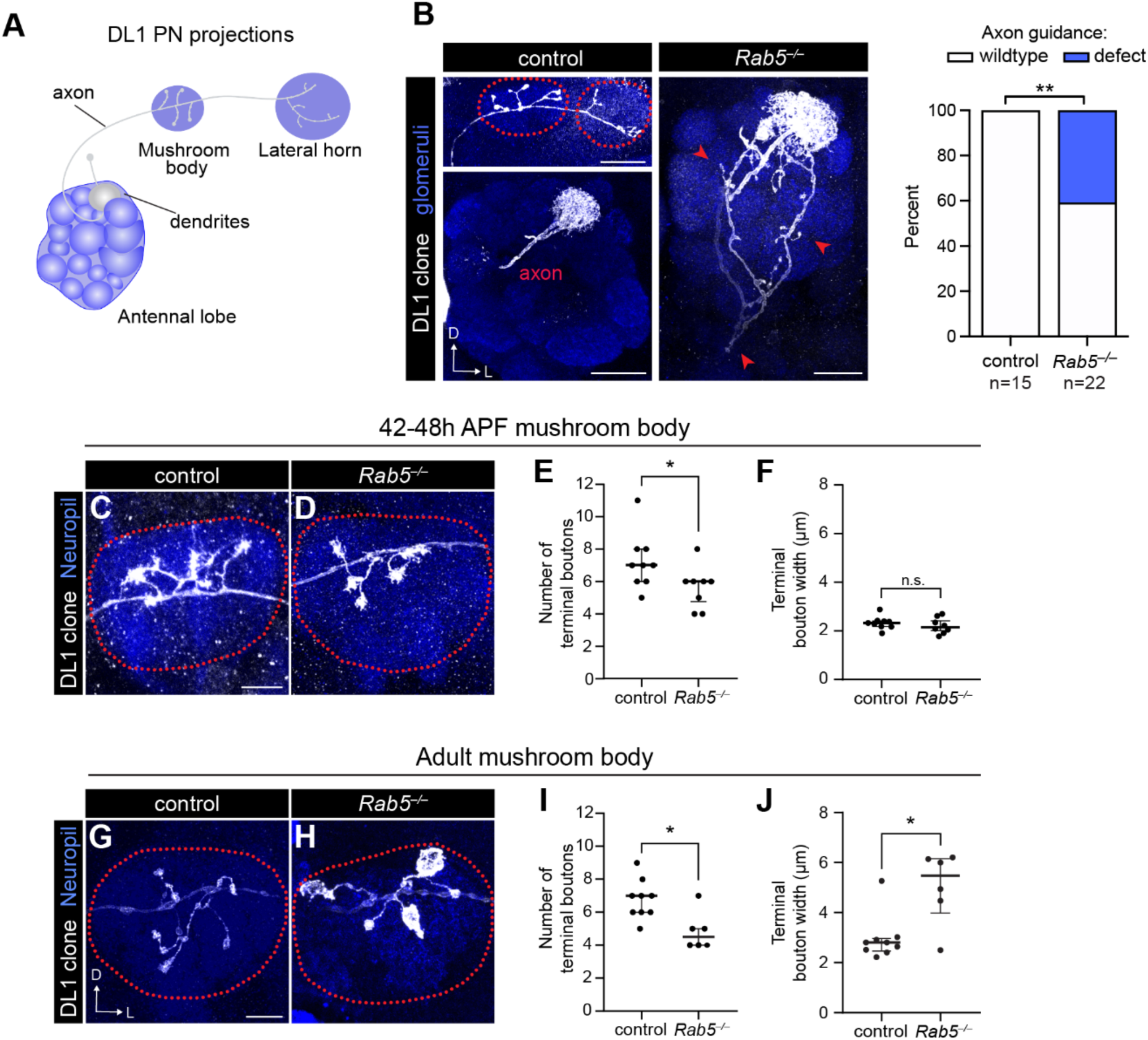
Rab5 is required for axon guidance and morphogenesis. (A) Schematic depicting DL1-PN dendrites innervating the DL1 glomerulus and axon projecting to the mushroom body and lateral horn. (B) Representative images of a control DL1 PN depicting its axon in the mushroom body and lateral horn (top left) and of the axon guidance defects observed in *Rab5* mutants where the axon does not project to higher order brain centers (middle) and quantification of the number of antennal lobes where this phenotype was observed (right). Red arrows denote DL1-PN axons terminating in ectopic regions. Fisher’s exact test was used to assess statistical significance. (C, D) Representative image of control (C) or *Rab5^−/−^* mutant (D) DL1-PN axons at the mushroom body at 48h APF. Dotted lines denote the border of the mushroom body. (E, F) Quantification of number of terminal boutons (E) and average terminal bouton width (F) in each DL1-PN mushroom body axon for controls (n=9) and *Rab5^−/−^* mutants (n=8) at 48h APF. Data are presented as medians (IQR); statistical significance was assessed using the Mann-Whitney U test. (G, H) Representative image of control (G) or *Rab5^−/−^* mutant (H) DL1-PN axons at the mushroom body at the adult stage. Dotted lines denote the border of the mushroom body. (I, J) Quantification of number of terminal boutons (I) and average terminal bouton width (J) in each DL1-PN mushroom body axon for controls (n=9) and *Rab5^−/−^*mutants (n=6) at the adult stage. Data are presented as medians (IQR). Mann-Scale bar, 20 µm (B); 10 µm (C, G).

We next examined the morphology of Rab5 mutant axons that successfully reached higher-order brain centers. After leaving the antennal lobe, DL1 axons first send branches that terminate in the mushroom body (Figure 3A), a neuropil critical for olfactory learning and memory (de Belle & Heisenberg, 1994; Heisenberg et al., 1985). Although the precise location of axon branches within this neuropil is stochastic, DL1 axons nonetheless display stereotyped morphology characterized by consistent bouton number, size, and branch number.

We first evaluated DL1 axon morphology during development (42–48h APF), as these axons were elaborating boutons and refining their morphology (Figure 3C, D). Since each axonal branch in the mushroom body terminates in a presynaptic bouton (Figure 3B, G), we quantified terminal boutons as a proxy for total axonal branch number. Compared to controls, *Rab5* mutant axons had fewer terminal boutons, representing a loss of higher order branching (Figure 3E). This reduction in axonal branching persisted into the adult mushroom body (Figure 3G–I) and was accompanied by a nearly two-fold increase in bouton size (Figure 3J), which was not observed during development (Figure 3F). These findings imply that mushroom body boutons retain the capacity to grow in later stages of development and suggest that Rab5 is required to restrain bouton expansion as the circuit matures. Together, these data indicate that Rab5 is required for axon branching while restricting bouton size in the mushroom body.

Beyond the mushroom body, PN axons innervate the lateral horn (Figure 3A). This second output region regulates innate olfactory responses (Heimbeck et al., 2001; Marin et al., 2002; Wong et al., 2002; Jefferis et al., 2007) and provides an opportunity to evaluate Rab5 function in a distinct axonal target witin the same neuron. DL1-PN projections to the lateral horn are stereotyped and exhibit a characteristic “L-shaped” branching pattern, composed of a shorter dorsal and longer lateral branch (Figure 3B), each of which contains relatively few higher-order branches. Compared to controls, *Rab5* mutant axons formed more secondary branches and boutons in the lateral horn (Figure 4A–D), indicating that Rab5 normally restrains branching and bouton elaboration this region. This contrasts its function in the mushroom body, where Rab5 promotes axon branching (Figure 3J, K), indicating that Rab5 has target-specific functions in axon terminal elaboration.

**Figure 4.**
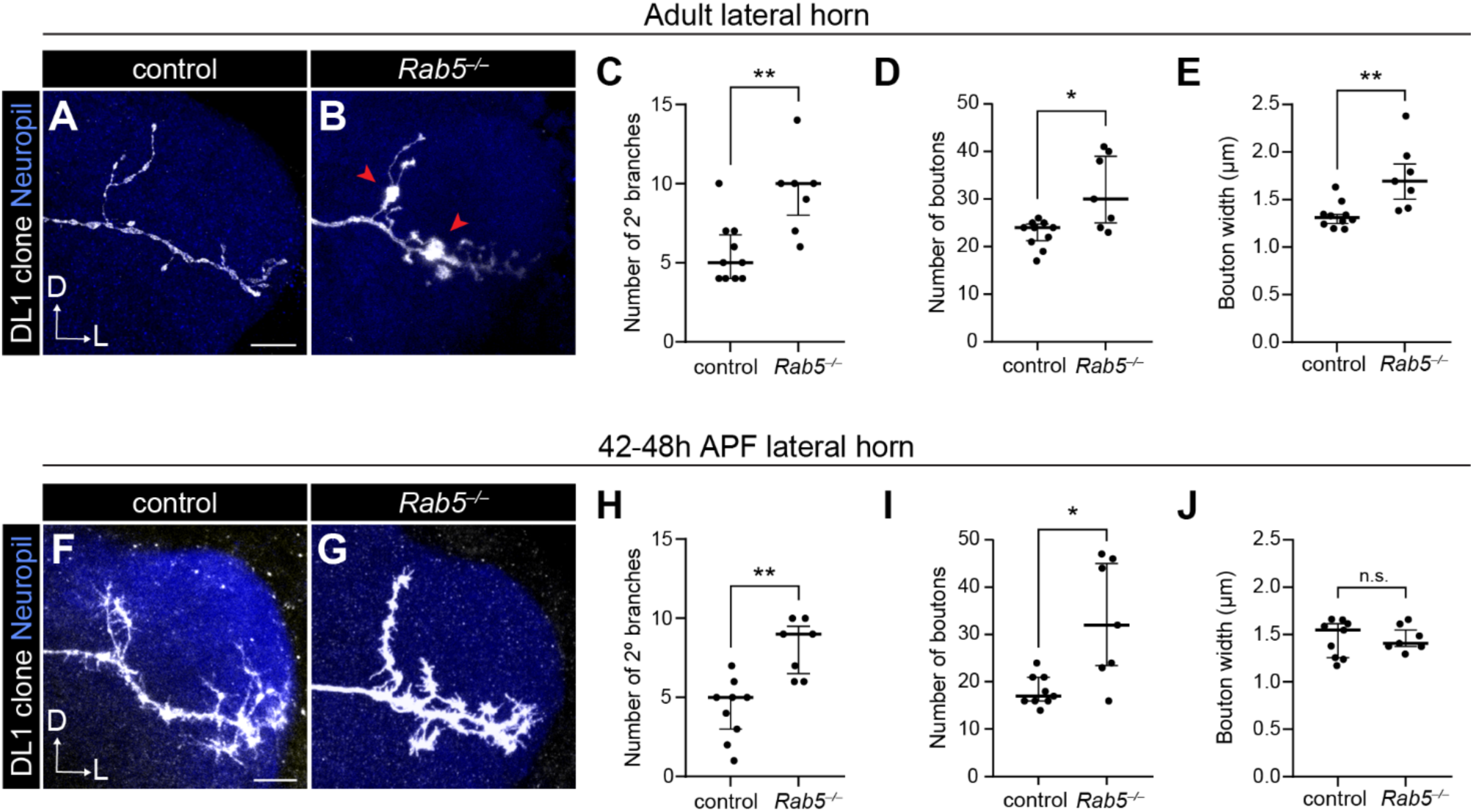
Rab5 restrains axonal morphogenesis in the lateral horn. (A, B) Representative image of a control (A) or *Rab5^−/−^* mutant (B) DL1-PN axon at the lateral horn at the adult stage. Red arrows denote enlarged boutons. (C–E) Quantification of number of secondary branches (C), number of boutons (D), and average bouton width (E) in each DL1-PN axon at the lateral horn for controls (n=10) and *Rab5^−/−^* mutants (n=7) at the adult stage. Data are presented as median (IQR); statistical significance was assessed using the Mann-Whitney U test. (F, G) Representative image of a control (F) or *Rab5^−/−^* mutant (G) DL1-PN axon at the lateral horn at 48h APF. Red arrows denote enlarged boutons. (H–J) Quantification of number of secondary branches (H), number of boutons (I), and average bouton width (J) in each DL1-PN axon at the lateral horn for controls (n=9) and *Rab5^−/−^* mutants (n=7) at 48h APF. Data are presented as median (IQR); statistical significance was assessed using the Mann-Whitney U test. Scale bar, 10 µm (A, F).

While most Rab5 mutant phenotypes were consistent between development (Figure 4F–I) and adulthood (Figure 4A–D), boutons in the lateral horn, like in the mushroom body, underwent substantial expansion after initial axon morphogenesis (Figure 4E, J). These findings indicate that PN bouton size remains plastic into later stages of development and that Rab5 acts as a critical brake on bouton expansion across distinct axonal targets. Altogether, these findings reveal that Rab5 can exert opposing effects on axon terminal arborization in different targets within the same neuron. Thus, highlighting that target-specific regulation of early endosomal trafficking is necessary to achieve the distinct axonal architectures.

### Developmental dynamics of early endosomes across PN compartments

To understand where Rab5 could be acting to carry out its functions, we mapped the distribution of early endosomes in PN dendrites and axons. Using MARCM, we expressed mCherry-2xFYVE, which predominantly accumulates on early endosomes (Gaullier et al., 1998; Gillooly et al., 2000), only in DL1-PNs at an early developmental timepoint (24–30h APF) and in adults. During development, there were relatively few early endosomes in PN dendrites, and they predominantly localized to more central dendrite regions and not the growing distal processes (Figure 5A). The adult dendritic arbor, on the other hand, was filled with many small early endosomes and larger clusters (Figure 5B). These findings indicate that the early endosome population expands in tandem with dendritic arbor growth.

**Figure 5.**
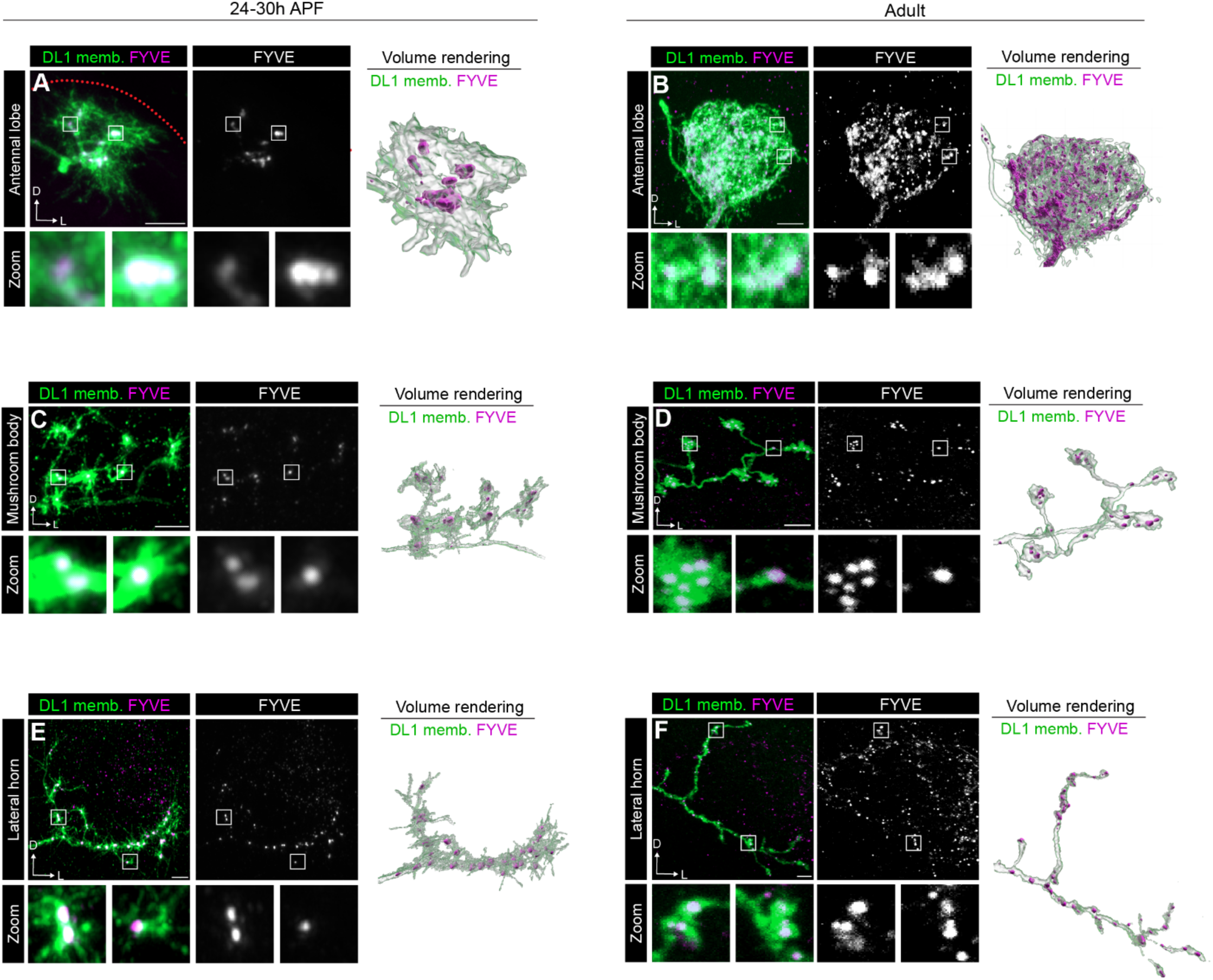
Developmental dynamics of early endosomes across neuronal compartments. (A) Airyscan super resolution images depicting the localization of FYVE-mCherry expressed in a DL1-PN single-cell clone (left), single-channel FYVE image (middle), and 3D volume rendering (right) at 24–30h APF. (B) Representative images depicting the localization of the early endosome marker FYVE-mCherry expressed in a DL1-PN single-cell clone (left), single-channel FYVE image (middle), and 3D volume rendering (right) at the adult stage. (C) Airyscan super resolution images depicting the mushroom body localization of the early endosome marker FYVE-mCherry expressed in a DL1-PN single-cell clone (left), single-channel FYVE image (middle), and 3D volume rendering (right) at 24–30h APF. (D) Representative images depicting the mushroom body localization of FYVE-mCherry expressed in a DL1-PN single-cell clone (left), single-channel FYVE image (middle), and 3D volume rendering (right) at the adult stage. (E) Airyscan super resolution images depicting the lateral horn localization of FYVE-mCherry expressed in a DL1-PN single-cell clone (left), single-channel FYVE image (middle), and 3D volume rendering (right) at 24–30h APF. (F) Representative images depicting the lateral horn localization of FYVE-mCherry expressed in a DL1-PN single-cell clone (left), single-channel FYVE image (middle), and 3D volume rendering (right) at the adult stage. Note that puncta outside of lateral horn axons are in other nearby cell types. Scale bar, 5 µm (A–F).

On the axon side, early endosomes accumulated in terminal boutons of both developing and adult mushroom body axons (Figure 5C, D), suggesting that terminal boutons are critical endosomal sorting sites. In the lateral horn, early endosomes were distributed throughout the axon but absent from actively growing processes during development (Figure 5E). In adult axons, this vesicle population was also found throughout the axon in both boutons and inter-bouton regions (Figure 5F). Altogether, these data indicate that early endosome distribution remains relatively stable between developing and adult stages in both axonal compartments, though their precise localization differs between the mushroom body and lateral horn.

### Rab7 is non-cell-autonomously required for DL1-PN dendrite targeting

Following entry into early endosomes, internalized cargo is directed toward one of two main destinations: lysosomes or recycling endosomes. However, the relative contributions of these distinct trafficking outcomes to the development of the same neuron *in vivo* remain largely opaque. To address this, we began by assaying Rab7 function in PN development. Rab7 predominantly localizes to late endosomes where it facilitates conversion of early endosomes to late endosomes (Rink et al., 2005), regulates late endosome fusion with lysosomes (Bucci et al., 2000; Vanlandingham & Ceresa, 2009; Xing et al., 2021), and promotes autophagosome maturation (Hyttinen et al., 2013).

The *Rab7* null allele is a GAL4 knock-in into the endogenous *Rab7* locus (Cherry et al., 2013) and is, therefore, incompatible with traditional MARCM approaches as they rely on the GAL4/UAS system to label cells (Lee & Luo, 1999; Wu & Luo, 2006). To circumvent this and analyze Rab7 function in PN development, we performed clonal analyses with a second binary expression system, the Q system (Potter et al., 2010; Potter & Luo, 2011), referred to as QMARCM (Potter et al., 2010). QMARCM is comparable to GH146-GAL4-based MARCM (Potter et al., 2010) but uses GH146-QF to drive QUAS transgenes carrying membrane markers or rescue transgenes.

We first quantified the number of adPN cell bodies and found that *Rab7* is dispensable for neurogenesis and/or neuronal survival (Figure 6-figure supplement 1A). Further, loss of *Rab7* resulted in mild dendrite targeting defects in neuroblast clones (Figure 6-figure supplement 1B, C, E, F). A few glomeruli (DL1, DC2, and DM6) had a reduction in innervation compared to controls (Figure 6-figure supplement 1C, E), but the rest were unchanged. To verify that Rab7 was responsible for these phenotypes, we generated a *QUAS-mCherry-Rab7* transgene and re-expressed it in adPN clones. Expression of mCherry-Rab7 in *Rab7* adPNs rescued the innervation defects to the DL1 and DM6 glomeruli and partially rescued targeting to DC2 (Figure 6-figure supplement 1D, E). In addition to the decrease in innervation, *Rab7* mutants exhibited minor mistargeting to non-adPN glomeruli (Figure 6-figure supplement 1F). While some of the mistargeting phenotypes could be rescued by re-expression of *mCherry-Rab7*, others could not (Figure 6-figure supplement 1D, F), indicating that either precise Rab7 levels are important for dendrite targeting or that *Rab7* loss in non-ad PNs could exert non-autonomous effects on neighboring neurons. This data suggests that Rab7 regulates dendrite targeting of a subset of PNs but is not broadly required for this process.

Does Rab7 have cell-autonomous roles in dendrite targeting? While neuroblast clones showed reduced innervation to the DL1 glomerulus (Figure 6—figure supplement 1A–C), *Rab7* mutant DL1-PN single-cell clones did not exhibit defects in dendrite targeting (Figure 6A, B). The failure to recapitulate the loss of innervation observed in neuroblast clones could indicate that (1) Rab7 is non-autonomously required for this process or (2) Rab7 perdurance may occur in the DL1-PN single-cell clones. Our initial findings that expressing Rab7-DN in most PNs caused dendrite targeting defects (Figure 1I) imply that Rab7 may be non-autonomously required for dendrite targeting. To circumvent potential RNA/protein perdurance in DL1-PN MARCM clones, we expressed Rab7-DN specifically in DL1-PNs using a DL1-PN-specific GAL4 driver and evaluated dendrite targeting. No mistargeting was observed when Rab7 function was impaired in DL1-PNs (Figure 6-figure supplement 2A, B). Thus, the DL1-PN dendrite mistargeting observed in adPN neuroblast clones is likely caused by disruption of Rab7 in neighboring PNs, implying that Rab7 is non-cell-autonomously required for dendrite targeting of DL1-PNs.

**Figure 6.**
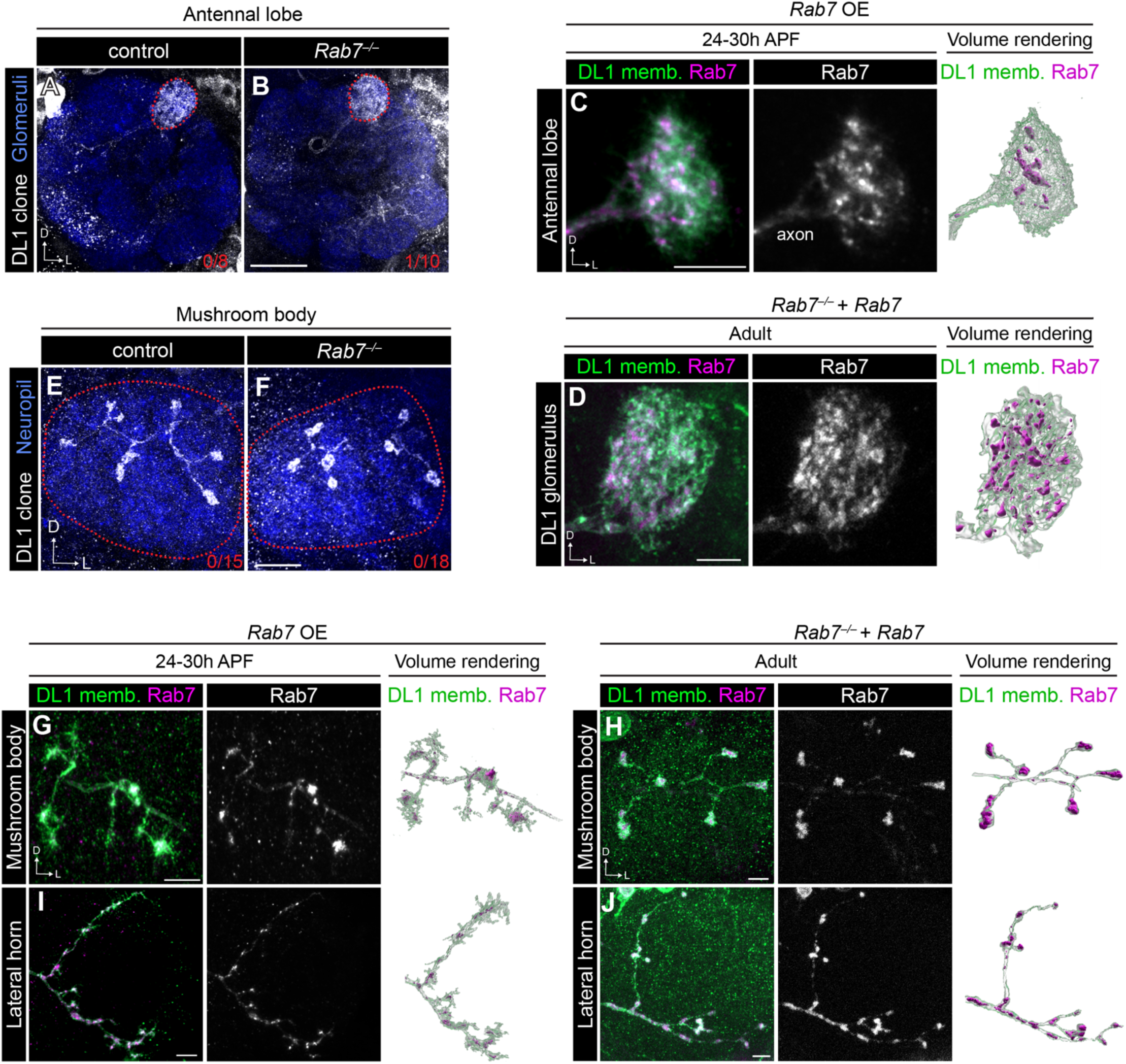
Rab7 plays a minor role in PN development. (A, B) Representative images of dendrite targeting of DL1-PN single-cell clones in control (A) and *Rab7^−/−^* mutant (B) antennal lobes. Red numbers in the right corner of images denote DL1-PN dendrite mistargeting phenotypic penetrance. (C) Airyscan super-resolution image depicting localization of mCherry-Rab7 in developing DL1-PN dendrites (left), single-channel image of Rab7 (middle), and 3D volume rendering of Rab7 distribution (right). (D) Representative image of localization of mCherry-Rab7 in a *Rab7^−/−^* mutant background in adult DL1-PN single-cell dendrites (left), single-channel image of Rab7 (middle), and 3D volume rendering of Rab7 distribution (right). (E, F) Representative images of mushroom body projections of control (E) and *Rab7^−/−^* mutant (F) DL1-PN single-cell clones. Red numbers in the right corner of images denote DL1-PN axon morphogenesis phenotypic penetrance. (G) Airyscan super-resolution image depicting localization of mCherry-Rab7 in a developing DL1-PN axon at the mushroom body (left), single-channel image of Rab7 (middle), and 3D volumes rendering of Rab7 distribution (right). (H) Representative image of localization of mCherry-Rab7 in a *Rab7^−/−^* mutant background in an adult DL1-PN axon at (I) Airyscan super-resolution image depicting localization of mCherry-Rab7 in a developing DL1-PN axon at the lateral horn (left), single-channel image of Rab7 (middle), and 3D volume rendering of Rab7 distribution (right). (J) Representative image of localization of mCherry-Rab7 in a *Rab7^−/−^* mutant background in an adult DL1-PN axon at the lateral horn (left), single-channel image of Rab7 (middle), and 3D volume rendering of Rab7 distribution (right). Scale bar, 20 µm (B); 10 µm (F); 5 µm (C, D, G–J).

Finally, we examined Rab7 localization in developing and adult dendrites. To evaluate Rab7 distribution at single-cell resolution, we re-expressed the mCherry-Rab7 transgene in *Rab7* mutant DL1-PN dendrites and evaluated its distribution. In adult dendrites, we found that Rab7 is punctate and present throughout the dendritic arbor (Figure 6D); however, we were unable to obtain reliable membrane GFP expression at 24-30h APF using the QMARCM system. We bypassed this limitation by generating and expressing a UAS version of mCherry-Rab7 in DL1-PNs using MARCM. During development, Rab7 was similarly present throughout the dendritic arbor (Figure 6C), suggesting that developing and adult dendrites may have comparable degradative needs.

### Rab7 does not play a major role in DL1-PN axon development

Does Rab7 play a role in axon development? To test this, we evaluated DL1-PN axon morphology in the mushroom body and lateral horn and found that *Rab7* mutant DL1-PN axons did not differ in morphology or branching compared to control DL1-PN axons in the mushroom body (Figure 6E, F) or lateral horn (Figure 6-figure supplement 1G). Consistently, expression of Rab7-DN in DL1-PNs did not alter any aspect of axonal morphology (Figure 6-figure supplement 2C–F). Thus, Rab7 may be dispensable for DL1-PN axon development.

Using the same genetic strategies as in dendrites, we evaluated Rab7 localization in mushroom body and lateral horn axons. In both developing and adult mushroom body axons, Rab7 appeared to be in large clusters (Figure 6G, H) within the boutons. In the lateral horn, Rab7 was abundant in both developing and adult projections (Figure 6I, J); however, unlike in the mushroom body, Rab7 puncta appeared larger in adult boutons than during developing ones, with many filled with prominent Rab7-positive clusters (Figure 6J). Overall, while Rab7 is expressed in developing neurons, Rab7-mediated degradation does not have a major role in regulating axon or dendrite development of DL1-PNs or could be acting redundantly with other degradative pathways.

### Rab11 has PN type-specific roles in dendrite targeting and innervation

Rab11 predominantly localizes to recycling endosomes and facilitates cargo recycling to the plasma membrane via this vesicular pool (Zulkefli et al., 2019). Thus, to understand how Rab11-mediated endosomal recycling contributes to circuit assembly, we started by testing Rab11’s role in PN dendrite targeting.

*Rab11* mutant adPNs exhibited decreased dendrite innervation into several adPN glomeruli as well as ectopic innervation into other non-adPN glomeruli (Figure 7A, B, D, E and Figure 7-figure supplement 1A), indicating that Rab11 regulates PN dendrite targeting. Loss of innervation to adPN glomeruli was not caused by impaired proliferation or increased cell death, as there was no difference in the number of adPNs in *Rab11* mutants compared to controls (Figure 7-figure supplement 1B). Re-expressing an mCherry-tagged Rab11 (Chen & He, 2022) in adPNs rescued the majority of the innervation defects (Figure 7C–E and Figure 7-figure supplement 1A), indicating that these defects are specific to *Rab11* loss. However, re-expression of Rab11 in *Rab11* adPN clones did cause some mistargeting to non-adPN glomeruli (Figure 7E and Figure 7-figure supplement 1A), suggesting that specific Rab11 levels may be important for the development of some PN types.

**Figure 7.**
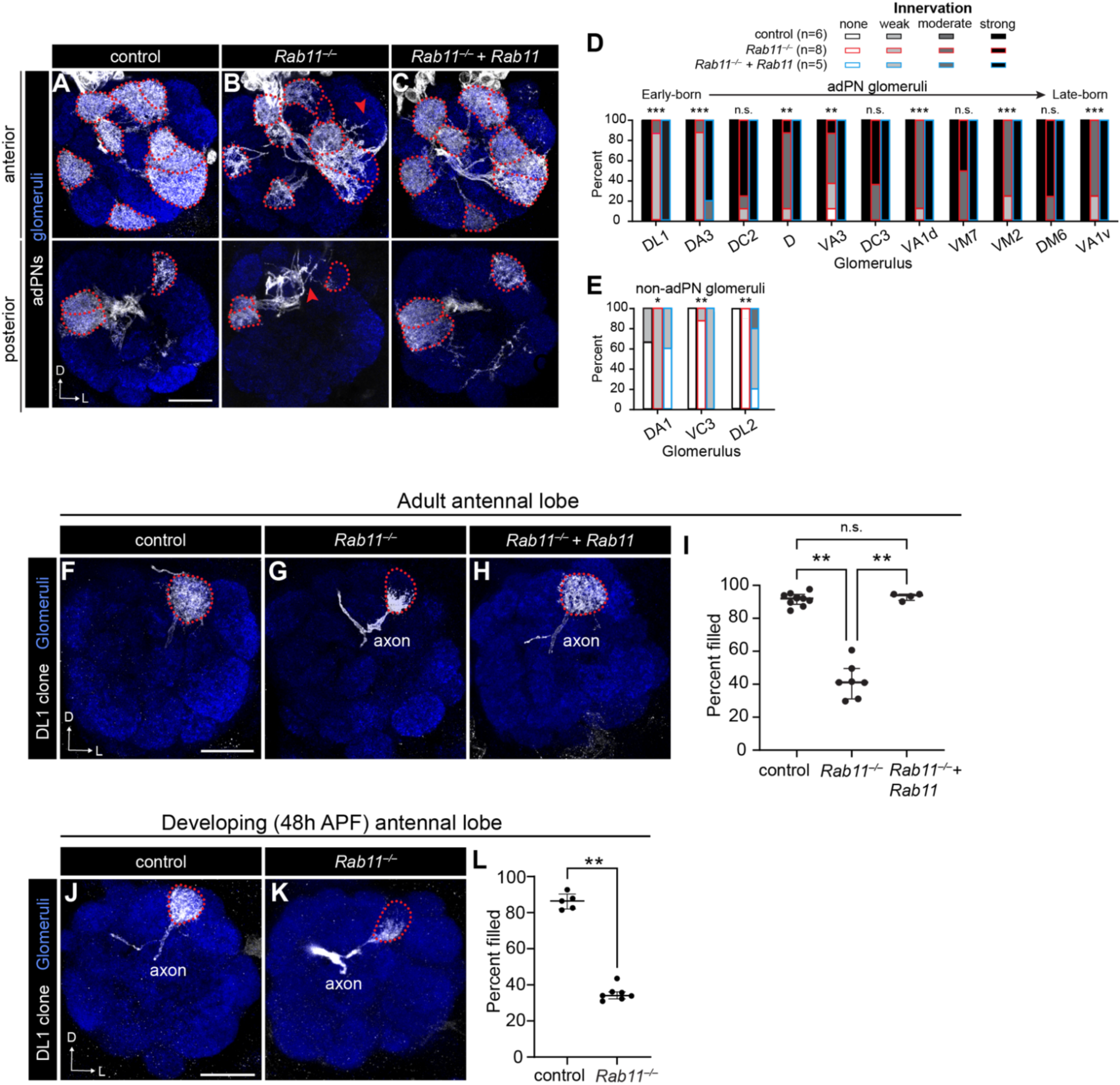
Rab11 has cell-type-specific roles in PN dendrite targeting and innervation. (A–C) Representative images of dendrite targeting of adPN neuroblast clones in control (A), *Rab11^−/−^* mutant (B), and *Rab11* rescue (C) antennal lobes. (D, E) Quantification of percent of antennal lobes with each category of dendrite innervation to adPN glomeruli (D) and glomeruli targeted by PNs other than larval-born adPNs (E). Fisher-Freeman-Halton test was used to assess statistical significance. See Table S3 for complete statistical analysis. (F–H) Representative images of control (F), *Rab11^−/−^* mutant (G), and *Rab11* rescue (H) DL1-PN single-cell clone dendrites. (I) Percent of the DL1 glomerulus filled by DL1-PN dendrites of controls (n=9), *Rab11^−/−^* mutants (n=7), and *Rab11* rescues (n=4). Data are presented as medians (IQR); statistical significance was assessed using a Kruskal-Wallis test followed by Dunn’s test post-hoc. (J, K) Representative images of control (J) and *Rab11^−/−^* mutant (K) DL1-PN single-cell clone dendrites at 42–48h APF. (L) Percent of the DL1 glomerulus filled by DL1-PN dendrites of controls (n=5) and *Rab11^−/−^* mutants (n=7) at 42–48h APF. Data are presented as medians (IQR); statistical significance was assessed using the Mann-Whitney U test. Scale bar, 20 µm (A, F); 15 µm (J).

The loss of innervation we observed in neuroblast clones could reflect either dendrite mistargeting to incorrect glomeruli or a failure of these dendrites to fully innervate within their appropriate target. To distinguish between these possibilities, we evaluated DL1-PN single-cell clones. DL1-PN clones did not exhibit dendrite mistargeting (Figure 7F, G, I). Rather, their dendrites failed to fully innervate the DL1 glomerulus and instead were concentrated at its ventromedial region, a phenotype that was fully rescued by cell-autonomous re-expression of *Rab11* (Figure 7H, I). Since defects in endosomal recycling are associated with neurodegeneration (Kiral et al., 2018), we probed if this innervation defect was the result of a failure to elaborate during development or retraction/degeneration in the adult stage. To answer this, we characterized DL1-PN dendrites during development. Similar to the adult phenotype, dendrites failed to fill the DL1 glomerulus, with only ∼40% coverage (Figure 7J–L). Together, these data indicate that Rab11 is required for dendrite elaboration.

Since *Rab11* mutant adPN neuroblast clones also exhibited ectopic targeting, we hypothesized that Rab11 could have cell-type-specific roles in dendrite targeting. Thus, we evaluated dendrite targeting in *Rab11* mutant DA1 or VA1d/DC3 neuroblast clones using *MZ19-GAL4-*based MARCM analysis. In neuroblast clones that contain labeled DA1-PNs, *Rab11* mutants exhibited reduced innervation into the DA1 glomerulus, along with ectopic targeting to mainly ventrolateral glomeruli (Figure 7-figure supplement 1C–E). Similarly, we observed a decrease in innervation into the VA1d and DC3 glomeruli and mistargeting to medial glomeruli in *Rab11* mutant neuroblast clones that contain labeled VA1d/DC3-PNs (Figure 7-figure supplement 1C–E). Importantly, *Rab11* mutants had a similar number of labeled cells as controls for both sets of clones (Figure 7-figure supplement 1F, G). Collectively, these data reveal that Rab11 has cell-type specific roles in PN dendrite targeting and innervation.

### Rab11 promotes axon stability and maturation

Next, we probed the role of Rab11 in DL1-PN axons. In the mushroom body, *Rab11* mutant DL1-PN axons had fewer secondary branches and terminal boutons along with increased terminal bouton size (Figure 8A, B, D-F), similar to *Rab5* mutants (Figure 3H, J). However, *Rab11* mutant terminal boutons also exhibited thin ectopic processes emanating from them (Figure 8B, G), reminiscent of the filopodia that are present during axon development (Figure 8 H, J). Most loss-of-function phenotypes could be rescued by re-expressing *Rab11* in DL1-PNs, indicating that Rab11 is cell-autonomously required to promote mushroom body axon branching and restrain bouton size (Figure 8C, D–G).

**Figure 8.**
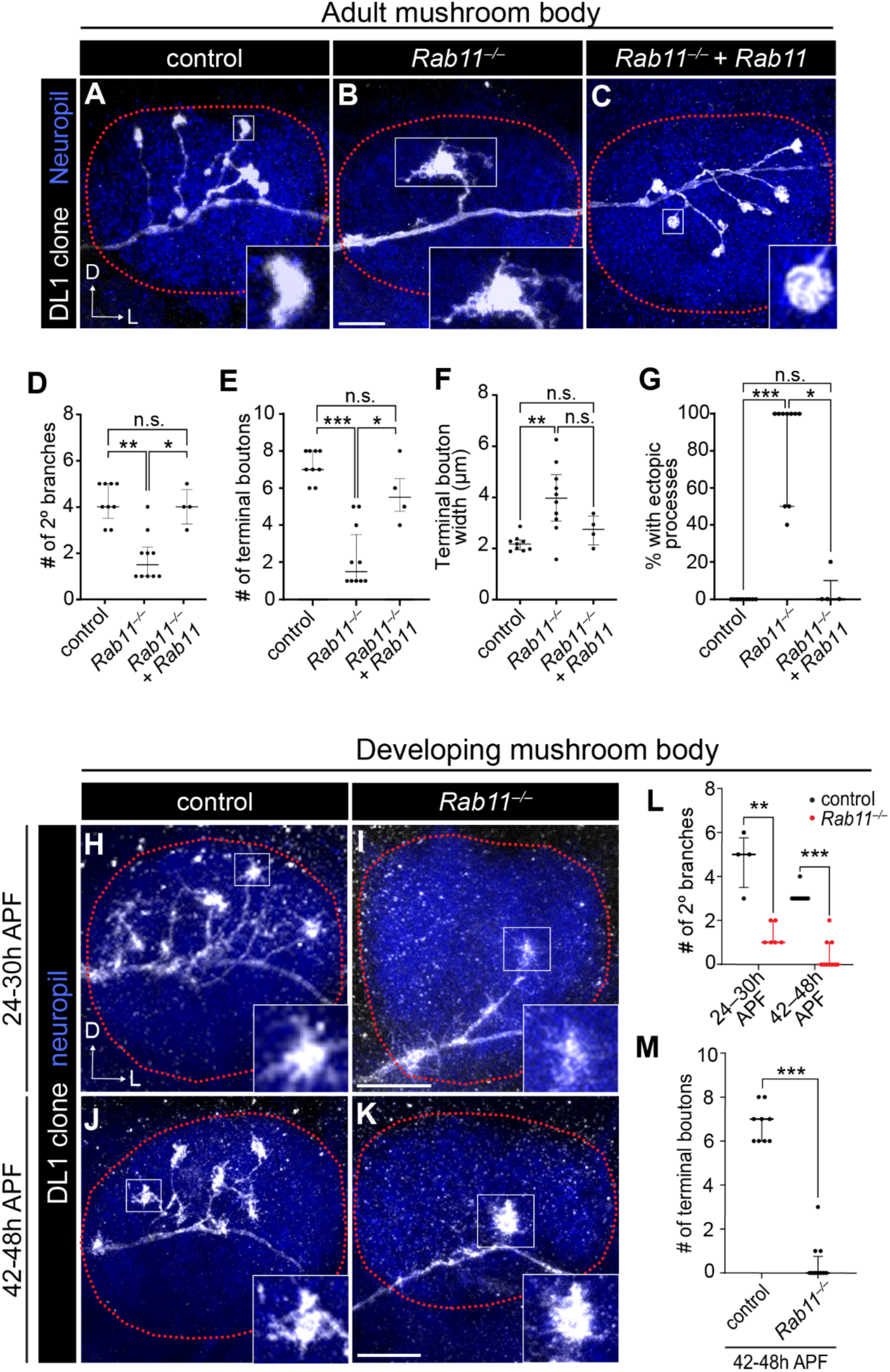
Rab11 promotes mushroom body axon maturation. (A–C) Representative images of adult control (A), *Rab11^−/−^* mutant (B), and *Rab11* rescue (C) DL1-PN axons in the mushroom body. (D–G) Quantification of the number of secondary branches (D), number of terminal boutons (E), average terminal bouton width (F), and percent of terminal boutons with ectopic processes (G) in each DL1-PN axon for controls (n=9), *Rab11^−/−^* mutants (n=10), and *Rab11* rescues (n=4). Data are presented as medians (IQR) and Kruskal-Wallis test followed by Dunn’s test post-hoc was used to assess statistical significance. (H–K) Representative images of developing control (H, J) and *Rab11^—/—^* mutant (I, K) DL1-PN axons in the mushroom body at 24-30h and 42-48h APF. (L) Quantification of the number of secondary branches in each DL1-PN axon at the mushroom body for 24-30h APF controls (n=4) and *Rab11^—/—^* mutants (n=6) and 42-48h APF controls (n=9) and Rab*11^—/—^* mutants (n=10). Data are presented as medians (IQR); statistical significance was assessed using the Mann-Whitney U test. (M) Quantification of the number of terminal boutons in each DL1-PN axon at the mushroom body for 42-48h APF controls (n=9) and Rab*11^—/—^* mutants (n=10). Data are presented as medians (IQR); statistical significance was assessed using the Mann-Whitney U test. Scale bar, 10 µm (B, I, K). Dotted lines denote the border of the mushroom body.

During development, DL1-PNs extend nascent axonal branches into the mushroom body around 12h APF which mature into defined collateral branches containing multiple boutons by approximately 50h APF (Jefferis et al., 2004; Zhu & Luo, 2004). The persistent filopodia present on *Rab11* mutant adult mushroom body axons implies that this GTPase promotes axon maturation. To evaluate if *Rab11* mutant axon phenotypes are the result of maturation defects, we examined these axons at two developmental timepoints: 24–30h APF and 42–48h APF. At the earlier timepoint, control axons were collateralized and contained nascent boutons with several filopodia-like processes (Figure 8H). However, *Rab11* mutant axons had fewer secondary branches and, therefore, a reduced number of terminal boutons (Figure 8I, L, M), indicating that during development they fail to elaborate the correct number of branches. By 42-48h APF, control axons had fewer filopodia-like structures and were beginning to resemble the mature terminal (Figure 8J). By this later timepoint, *Rab11* mutants had lost most of their secondary branches and often contained a single branch or large bouton, similar to phenotypes observed in the adult stage (Figure 8B, K–M) indicating that Rab11 is required for both axon branch formation and stability. Together, Rab11 is required for mushroom body axon maturation and stabilization.

Does Rab11 play similar roles in the lateral horn? Indeed, loss of *Rab11* resulted in multiple morphological defects in these targets in the adult stage. ∼50% of *Rab11* mutant axons lacked either the dorsal or lateral branch (Figure 9A, B [left], D). Among those with a dorsal collateral, ∼50% exhibited overextension of this process, and some had a second dorsal branch (Figure 9B [left], E), which is occasionally observed in pupal stages but does not persist in adults (Jefferis et al., 2004). In addition to branching defects, *Rab11* mutant axons extended many short filopodia-like processes (Figure 9B) similar to the nascent processes we observed in the mushroom body (Figure 8). We quantified this by measuring the number of endpoints and average length of each process emanating from the dorsal or lateral branches and found that *Rab11* mutant axons contained a significantly higher number endpoints and shorter process lengths than controls (Figure 9F, G). Many of these phenotypes were rescued by re-expressing *Rab11* in DL1 PNs (Figure 9C, D–G), indicating that Rab11 cell-autonomously promotes lateral horn axon development.

**Figure 9.**
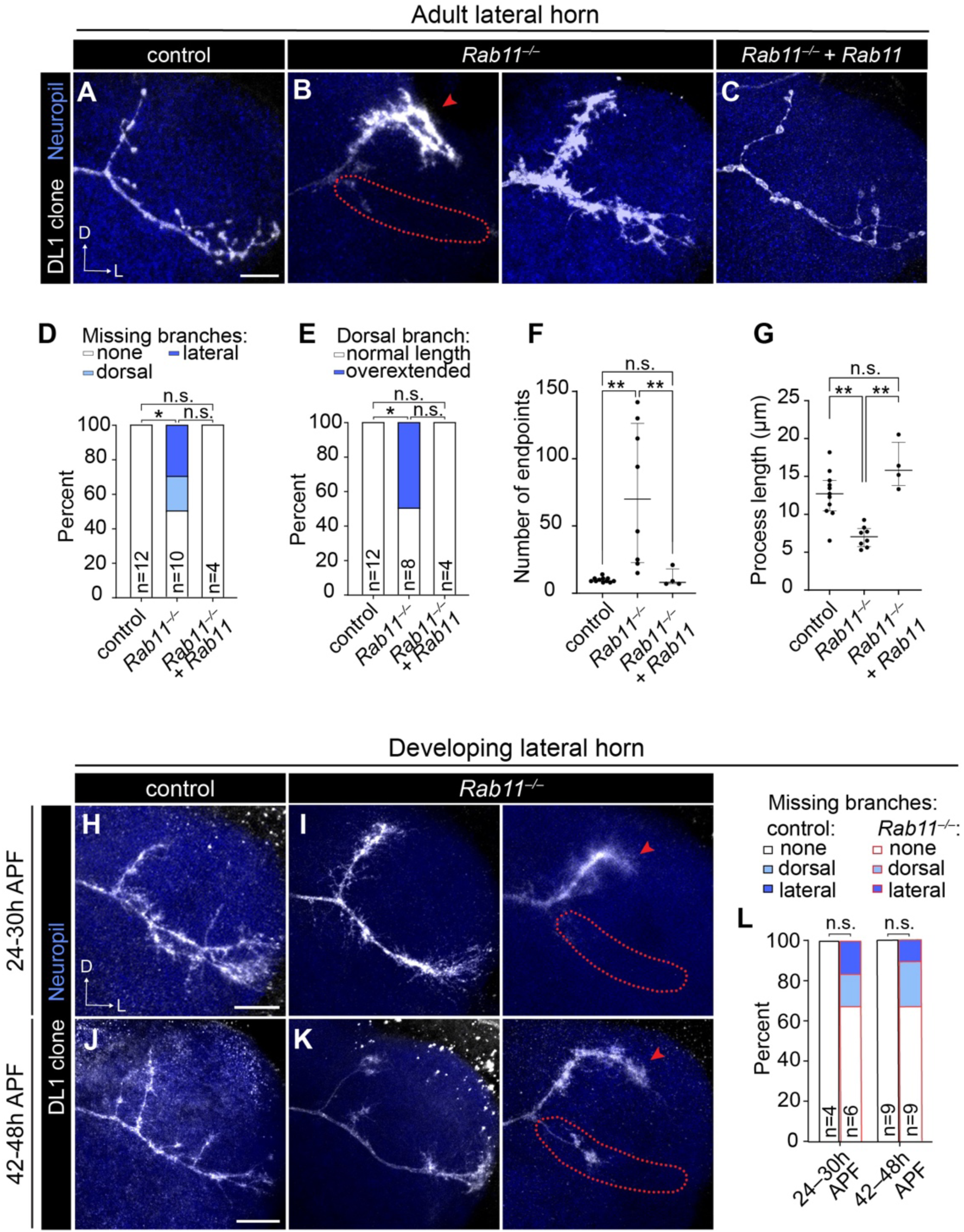
Rab11 is required for the maturation of lateral horn axons. (A–C) Representative images of adult control (A), *Rab11^−/−^* mutant (B), and *Rab11* rescue (C) DL1-PN axons at the lateral horn. (D, E) Quantification of the percent of axons missing a main branch (D) or percent with an overextended dorsal branch (E). Statistical significance was assessed using the Fisher-Freeman-Halton test, followed by pairwise Fisher’s exact tests post-hoc.(F, G) Quantification of number of endpoints (F) or average process length (excluding main branches) (G) of each DL1-PN axon in the lateral horn of controls (n=11), *Rab11^−/−^*mutants (n=8), and *Rab11* rescues (n=4). Data are presented as medians (IQR). Kruskal-Wallis test followed by Dunn’s test post-hoc was used to assess statistical significance. Some measures did not reach statistical significance due to a low sample number inherent with MARCM analysis. (H–K) Representative images of developing control and Rab*11^—/—^* mutant DL1-PN axons in the lateral horn at 24–30h and 42–48h APF. (L) Quantification of the percentage of lateral horn axons that have missing branches in each genotype at indicated developmental timepoints. Fisher’s exact test was used to assess statistical significance Scale bar, 10 µm. Dotted lines denote the border of the loss of lateral branches. Arrow heads indicate overextension of the dorsal branch.

We next examined developing lateral horn axons to determine if the branching defects in *Rab11* mutant axons resulted from a failure to form branches or from branch retraction. The lateral branch extends first (at ∼18h APF) followed by the appearance of the dorsal collateral between 24–30h APF (Jefferis et al., 2004). By 42h APF, a more mature axonal morphology emerges, characterized by the loss of filopodia-like projections and the appearance of rounded boutons (Figure 9H vs. J). Developing *Rab11* mutant axons never lost these filopodia and additionally lacked either a dorsal or lateral branch (Figure 9I, K [right]), similar to the phenotypes at the adult stage. The proportion of *Rab11* mutants missing the lateral branch at the adult stage was approximately 2.5-fold higher than at either developmental stage, whereas the proportion of mutants missing the dorsal branch remained similar across all stages (Figure 9D, L). These findings suggest that Rab11 plays distinct roles in lateral horn axon branching by promoting dorsal branch formation and preventing lateral branch retraction. Further, they highlight that Rab GTPases can exert target-specific effects within the same neuron. Collectively, our data demonstrates that Rab11-mediated recycling is required for axon maturation and morphological stability.

### Developmental dynamics of Rab11 in PNs

Since Rab11 was broadly required for PN development, we sought to visualize its distribution to gain insight into where it might be acting. Because re-expression of mCherry-Rab11 rescued most *Rab11* mutant phenotypes, we leveraged this genetic background to visualize Rab11 localization in developing and adult DL1-PN clones. During development, Rab11 was largely excluded from the distal dendritic processes and was instead concentrated at branch points, suggesting that recycling endosomes are enriched at central dendritic hubs during arborization (Figure 10A). In adult neurons, Rab11 was distributed throughout the arbor, including in distal processes (Figure 10B). The increased number and density of Rab11-positive compartments in adult dendrites suggests that as dendrites grow and elaborate there is a concomitant expansion of the recycling endosome network to support their functional demands.

**Figure 10.**
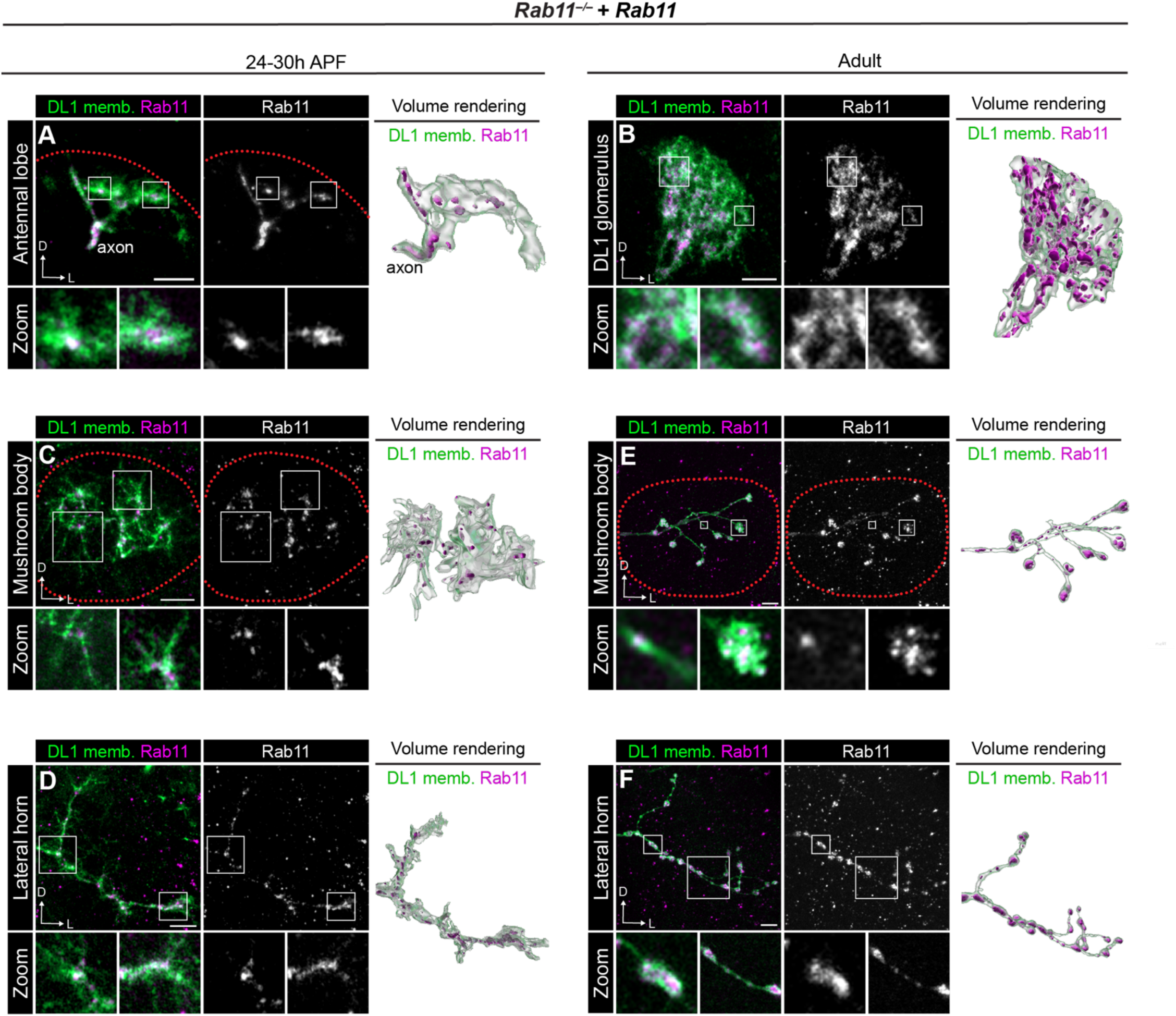
Developmental dynamics of Rab11 distribution in PNs. (A) Airyscan super resolution images depicting the localization of mCherry-Rab11 in a *Rab11^−/−^* mutant background in a DL1-PN single-cell clone dendrites (left), single-channel Rab11 image (middle), and 3D volume rendering (right) at 24–30h APF. Zoom panels are single optical sections of areas indicated by white boxes. (B) Representative images depicting mCherry-Rab11 localization in a *Rab11^−/−^* mutant background in a DL1-PN single-cell clone dendrites (left), single-channel Rab11 image (middle), and 3D volume rendering (right) at the adult stage. Zoom panels are single optical sections of areas indicated by white boxes. (C, D) Airyscan super resolution images depicting mCherry-Rab11 localization at the mushroom body (C) and lateral horn (D) in a *Rab11^−/−^* mutant background in a DL1-PN single-cell clone (left), single-channel Rab11 image (middle), and 3D volume rendering (right) at 24–30h APF. Zoom panels are single optical sections of areas indicated by white boxes. (E, F) Representative images depicting mCherry-Rab11 localization at the mushroom body (E) and lateral horn (F) in a *Rab11^−/−^* mutant background in a DL1-PN single cell clone (left), single-channel Rab11 image (middle), and 3D volume rendering (right) in the adult stage. Zoom panels are single optical sections of areas indicated by white boxes. Scale bar, 5 µm (A–F).

We next examined Rab11 localization in axons. Similar to developing dendrites, Rab11 was enriched at axonal branchpoints in both mushroom body and lateral horn axons, with reduced localization to actively growing processes (Figure 10C, D). By the adult stage, Rab11 accumulated in boutons with fewer smaller puncta in the inter-bouton regions (Figure 10E, F). Rab11 enrichment in mature boutons of both lateral horn and mushroom body axons could reflect its well-defined role in synaptic vesicle recycling (Ivanova & Cousin, 2022). Altogether, these observations indicate that Rab11 localization is dynamically regulated across development likely reflecting changes in recycling needs or cargo distribution across time.

## DISCUSSION

Building functional neural circuits depends on the coordination of diverse cellular processes each governed by tightly regulated protein trafficking, signaling, and turnover. *Drosophila* PNs provided a powerful model system to systematically evaluate the neurodevelopmental functions of individual endosome-associated Rabs within the same cell types *in vivo.* Through this approach, we found that even within a single neuron, distinct post-endocytic sorting events regulate different aspects of development and, in some cases, act in a compartment-specific manner (Figure 11). Below we discuss how studying membrane trafficking events *in vivo* can expand our understanding of neuronal development and cell biology.

**Figure 11.**
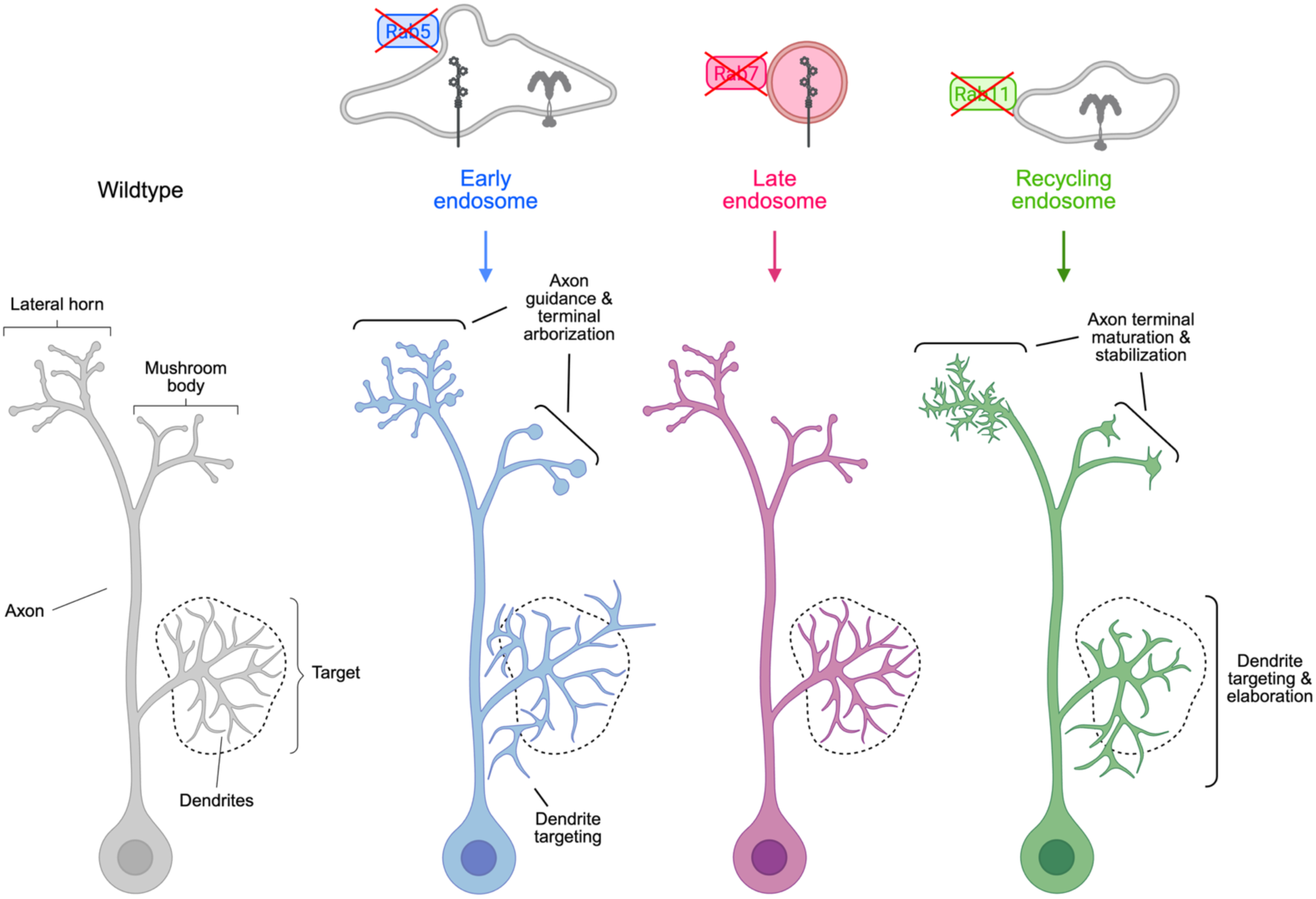
Endosome-associated Rab GTPases regulate distinct aspects of neuronal morphogenesis and circuit assembly. Summary model of the roles of endosome-associated Rab GTPases in PN development.

Among the Rabs we tested, Rab5 regulated the broadest range of developmental events—likely reflecting its role in establishing the initial sorting hub from which many internalized proteins are routed. In contrast, Rab7 and Rab11 had more specialized roles with Rab11 directing a larger set of developmental processes than Rab7. Notably, Rab11 and Rab5 regulated largely non-overlapping processes, except for promoting axon branching and restraining bouton size in the mushroom body. These distinctions suggest that additional Rab-mediated trafficking routes downstream of early endosomes are also critical for circuit assembly.

Despite extensive study, the role of Rab5 in neuritogenesis and growth remains debated, with some reports indicating that it inhibits these processes and others finding the opposite (Villarroel-Campos et al., 2016). These inconsistencies may stem from the reliance on overexpression systems, dominant-negative constructs, or *in vitro* models. Our clonal analysis approach enabled us to directly probe the role of Rab5 in circuit assembly. Our findings, which extend those from a prior study (Sakuma et al., 2014), indicate that neurites still form and extend in the absence of Rab5, but both axons and dendrites fail to properly navigate to their appropriate targets. This is consistent with another report that found Rab5 is required for axon targeting of cortical neurons (Wu et al., 2014). Extending these insights, we defined additional roles for Rab5 and found that it is required to restrain PN axon branching and bouton size—processes that are distinct from initial neurite growth and guidance. We also showed that early endosomes are distributed in nascent boutons where they can exact their developmental functions. Collectively, our data highlight the pleiotropic roles of Rab5 in multiple neurodevelopmental processes.

Although Rab7 is the primary regulator of late endosome-lysosome fusion (Bucci et al., 2000; Vanlandingham & Ceresa, 2009; Xing et al., 2021), our data suggest that it plays a minor role in PN development—consistent with prior studies of dendritic (Harish et al., 2019) and axonal (Ponomareva et al., 2016) branching. Specifically, expression of Rab7-DN in DL1-PNs and single-cell *Rab7* mutant MARCM clones both failed to produce dendrite targeting or axonal phenotypes. Some residual Rab7 activity may remain in both manipulations due to RNA/protein perdurance in MARCM clones or incomplete dominant negative suppression. Nevertheless, our findings suggest that Rab7-mediated degradation is not a major driver of DL1-PN development, though whether this reflects true dispensability or simply a lower threshold requirement for Rab7 activity remains an open question.

Rab7 did have non-autonomous effects on dendrite targeting for the PN types we examined. This is in line with findings from embryonic development where Rab7 was shown to non-autonomously regulate receptor signaling during gastrulation (Kawamura et al., 2020). Perturbing Rab7 or endosomal acidification has been shown to cause a buildup of early endosomes (Girard et al., 2014; Lelouvier & Puertollano, 2011; Martina et al., 2009), raising the possibility that in some PNs impaired degradation of receptors could prolong endosomal signaling and lead to non-autonomous effects on nearby neurons.

A possible explanation for the absence of cell-autonomous phenotypes is compensation by other degradative Rabs. While Rab9 and Rab2 associate with degradative compartments, Rab9 is not expressed in developing PNs (Figure 1-figure supplement 1), making it unlikely to compensate for Rab7 loss. Rab2 is expressed in PNs but primarily functions in autophagosome-lysosome fusion (Ding et al., 2019) and trafficking of lysosomes and their associated proteins toward late endosomes (Lund et al., 2018), rather than directing internalized cargo towards degradative compartments. Thus, while Rab2 is associated with the degradative pathway it does not have overlapping functions with Rab7. Collectively, our findings imply that Rab7-mediated degradation plays a limited role in DL1-PN development, provided that upstream trafficking events adequately remove receptors from the plasma membrane and direct them towards endosomal compartments.

Rab11, on the other hand, was important for axonal maturation, particularly in the formation and stabilization of boutons and branches. Strikingly, even within the same neuron, Rab11 regulated distinct aspects of axon development: in the mushroom body, Rab11 promoted axonal branching, whereas in the lateral horn, it was critical for the formation of certain branches and the stabilization of others. Given Rab11’s well-established role in endosomal recycling, these phenotypes are consistent with a failure to deliver adhesion and signaling receptors back to the plasma membrane, disrupting maturation of nascent axonal processes. Further, the target-specific effects of Rab11 (in mushroom body vs. lateral horn projections) suggests that distinct sets of cargos may be recycled in different axonal regions (or that the same cargo could have distinct functions in different axonal targets) to drive local developmental processes.

Underscoring the specificity of endocytic recycling, Rab11 regulated distinct aspects of dendrite development across PN types. In DL1-PNs, Rab11-mediated recycling was critical for dendrites to fully innervate their target glomerulus, whereas this GTPase promoted dendrite targeting in DA1-PNs and VA1d/DC3-PNs. These findings not only extend the role for Rab11 beyond just dendrite branching (Takano et al., 2014) but also suggests that it could recycle a unique set of cargoes in each cell type. Supporting this idea, the Rab11-dependent DL1-PN dendrite phenotype, but not the VA1d-PN phenotype, resembles that observed upon loss of the cell adhesion protein Dscam, which normally promotes dendritic self-avoidance (Zhu et al., 2006). These data raise the possibility endocytic recycling regulates self-recognition events critical for dendrite elaboration, though further studies are needed to directly test this. Thus, our findings highlight that endosomal recycling can exert cell type- and compartment-specific control over neuronal development. Defining the cargos regulated by Rab11-mediated trafficking will be essential for understanding how this pathway sculpts connectivity. Altogether, our work reveals how neurons harness multiple endocytic routes, in a compartment- and cell type–specific manner, to control neuronal morphogenesis and circuit formation.

## ACKNOWLEDGEMENTS

We are grateful to C. Taylor and J. Kalai for their helpful feedback on this manuscript. We thank members of the Luo lab and E. Theisen, L. Jane, and Z. Cook for support, insight, and feedback on this study. B. He (Dartmouth College), R. Hiesinger (Freie Universität Berlin), Addgene, Bloomington *Drosophila* Stock Center, Vienna *Drosophila* Stock Center, and Best Gene, provided critical reagents. We appreciate the administrative assistance from M. Molacavage. C.N.M. was a HHMI fellow of the Damon Runyon Cancer Research Foundation (DR-2390-20). L.L is a HHMI investigator. This work was supported by the National Institutes of Health (R01-DC005982 to L.L. and K99-DC021195 to C.N.M.).

## AUTHOR CONTRIBUTIONS

C.N.M. conceived this project. K.X.D. and C.N.M. designed and completed the experiments with assistance from H.J. and Y.Z. D.J.L. generated new fly lines used in this study. K.X.D. analyzed data and generated figures. C.N.M. wrote the manuscript with input from K.X.D. and L.L. and all other coauthors. C.N.M. and L.L. supervised the work.

## DECLARATION OF INTERESTS

The authors declare no competing interests.

## MATERIALS AND METHODS

### Materials availability

We have deposited newly generated flies in the Bloomington Drosophila Stock Center. All other unique reagents generated in this study are available from the corresponding author

### Key Resources Table

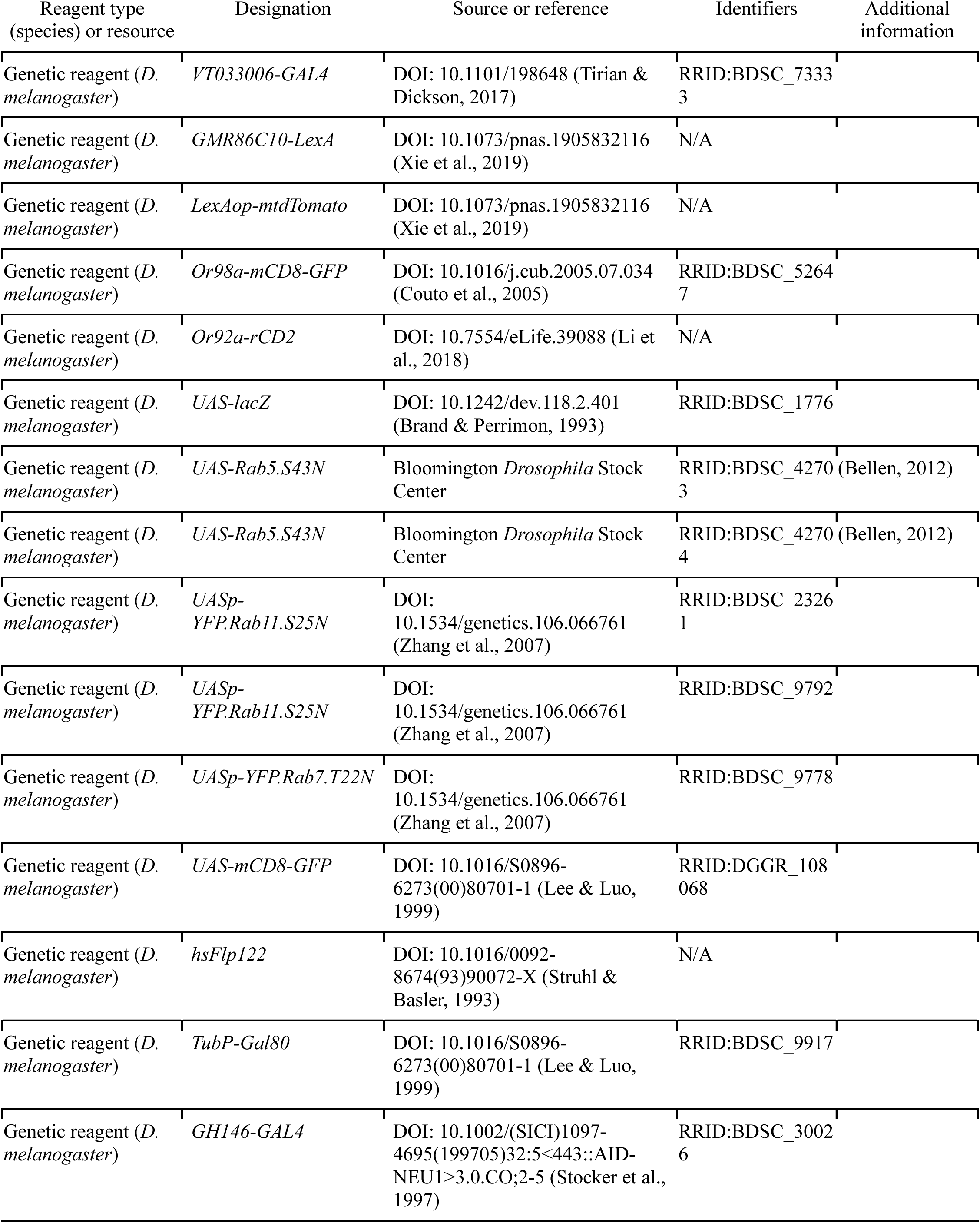
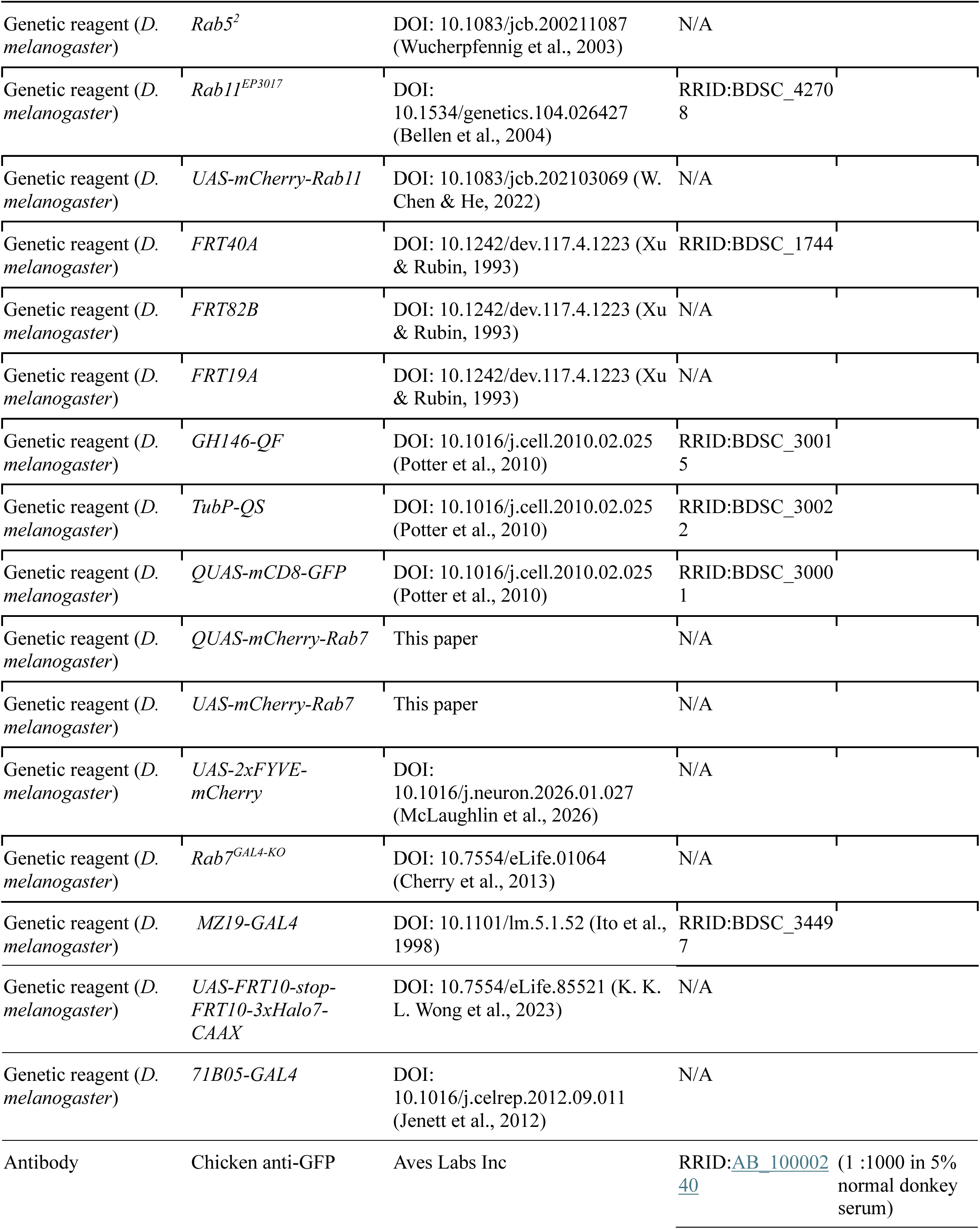
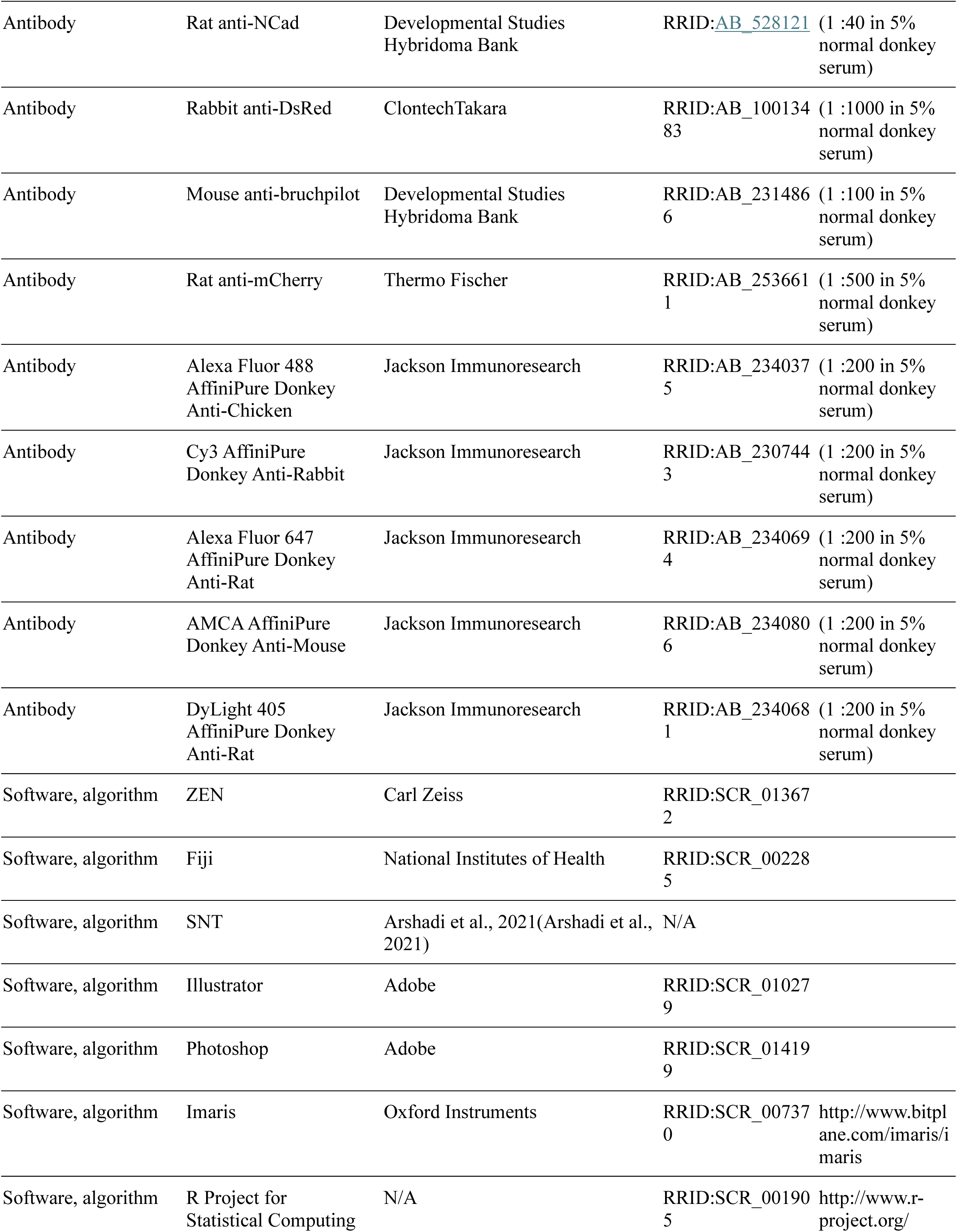

### *Drosophila* stocks and husbandry

Flies were maintained on standard cornmeal media with a 12 hr light-dark cycle at 25°C, except for overexpression crosses which were raised at 29°C. The strains used are described in the Key Resources Table and complete genotypes for flies used in each figure are in Table S1.

### Generation of UAS/QUAS constructs and transgenic flies

The *UAS-mCherry-Rab7* and *QUAS-mCherry-Rab7* constructs were synthesized and cloned by Twist Biosciences into a 10x pUASt-attB vector or 10x QUAS-attB vector. The mCherry tags are N-terminal to preserve Rab function. The construct was validated by full-length plasmid sequencing and injected into embryos with an attP40 insertion site. G0 flies were crossed to a white– balancer and all white+ progeny were individually balanced. Flies were injected in-house using standard microinjection methods.

### Immunofluorescence staining and confocal microscopy

Fly brains were dissected according to a previously published protocol (J. S. Wu & Luo, 2006a). In brief, brains were dissected in PBS, transferred to a tube containing 4% paraformaldehyde in PBST (0.3% Triton X-100), and fixed for 20 min while nutating at RT. Following fixation, brains were washed 3 times for 20 minutes in PBST and blocked for at least 30 min in PBST + 5% normal donkey serum. The following antibodies were used: rat anti-Ncad (Developmental Studies Hybridoma Bank [DSHB]; 1:40), chicken anti-GFP (Aves Labs; 1:1000), rabbit anti-dsRed (Takara Bio; 1:1000), mouse anti-bruchpilot (DSHB; 1:100), mouse anti-mCherry (ThermoFisher Scientific; 1:1000) and incubated with brains in block buffer overnight at 4°C while nutating. Brains were subsequently washed three times for 20 min in PBST and incubated in secondary antibodies (Alexa Fluor 488; Alexa Fluor 564; Alexa Fluor 647; 1:200) overnight at 4°C while nutating. Brains were again washed three times for 20 min in PBST, transferred to SlowFade antifade reagent (ThermoFisher) and stored at 4°C prior to mounting.

### Image acquisition and processing

Images were obtained on a Zeiss LSM900 laser-scanning confocal microscope (Carl Zeiss) using either a 40x oil immersion objective (dendrite targeting and axon morphology experiments) or a 63x oil immersion objective (Airyscan experiments). 16-bit z-stacks for dendrite targeting and axon morphology experiments were acquired at 1 µm intervals at a resolution of 1024 × 1024. Airyscan images were taken at software optimized resolution and intervals. Brightness and contrast adjustments as well as image cropping was done using Photoshop or Illustrator (Adobe).

### Dominant negative screen

Virgin *Drosophila* females with the genotype *VT033006-GAL4, GMR86C10-LexA>LexAop-mtdTomato, Or98a-mCD8::GFP, Or92a-CD2* were crossed with *UAS-RabX-DN* males to express GDP-locked dominant negative Rabs in *VT-GAL4* PNs, and the progeny were kept at 25°C for 2–5 days following egg laying and then transferred to 29°C to enhance transgene expression. Brains were dissected, processed, and imaged as described above. See Table S1 for complete genotypes.

For this analysis, we identified glomeruli using NCad labeling (based on the stereotypy of their size, shape, and positions). VM5d/v PNs were monitored through the expression of mtdTomato using the *GMR86C10-LexA* driver, and dendrite targeting was categorized by the presence or absence of mistargeting to ectopic glomeruli. Dendrite targeting analysis was performed blinded to genotype when possible. Fisher’s exact test was performed on mistargeting frequencies to determine statistical significance compared to controls. P-values were adjusted using the Benjamini-Hochberg procedure.

### MARCM-based clonal analyses

Clonal analyses using mosaic analysis with a repressible cell marker (MARCM) and Q-MARCM (MARCM using the Q system) have been previously described (Potter & Luo, 2011; J. S. Wu & Luo, 2006b). Each fly contains a *hsFLP122* recombinase, *GH146-GAL4* (PN GAL4) or *GH146-QF* (PN QF), *TubP*-*Gal80* or *TubP*-*QS*, *UAS-mCD8-GFP* or *QUAS-mCD8-GFP*, the desired *FRT*, and either wild-type or a mutant allele distal to the *FRT* site; flies for the Rab11 rescue experiments also included a *UAS-mCherry-Rab11*, and flies for the Rab7 rescue experiments included a *QUAS-mCherry-Rab7* (see Table S1 for complete genotypes). To generate adPN neuroblast, DL1 single-cell, DA1 neuroblast, and VA1d/DC3 neuroblast clones, flies were heat shocked for 1 hour at 37 °C at 0-24h after larval hatching. To generate smaller neuroblast clones with projections to the DM6 glomerulus, flies were heat shocked for 1 hour at 37 °C at 48-72h after larval hatching. Brains were dissected and processed as described above. We identified glomeruli using NCad labeling and categorized the extent of innervation into each glomerulus (not innervated; weakly innervated; moderately innervated; strongly innervated). Analysis was performed blinded to experimental manipulation. For each glomerulus, we calculated the frequency of each type of innervation and plotted the results as stacked bar charts. Fisher’s exact test was performed on innervation frequencies in each glomerulus, using counts for wildtype levels of innervation (strong if innervation is expected and none if innervation is not expected in wildtype) vs. non-wildtype levels of innervation, to determine statistical significance compared to controls. P-values were adjusted using the Benjamini-Hochberg procedure.

### Image analysis and quantification

Microscopy images were processed and analyzed using ImageJ tools. Dendrite glomerular innervation was measured using the area measuring tool where the area of each DL1 glomerulus was measured using NCad staining and compared to the area of GFP-positive dendrites within that glomerulus. Bouton diameter was measured by drawing a line across the widest part of each bouton and measuring the length using the length measuring tool. Axon branches were traced and measured using Semi-automated Tracing on GFP-positive axons in the Simple Neurite Tracer (SNT) plugin (Arshadi et al., 2021) to quantify branch numbers and lengths. All analyses were performed blinded to genotype.

Volume renderings were created using Imaris10 (Oxford Instruments); Airyscan super-resolution images (Carl Zeiss) were imported, and the Surfaces tool was used to model the membrane of GFP-positive dendrites and axons as well as mask the puncta channel. The Surfaces tool was used to model puncta with the masked puncta channel, and the Spots tool was used to quantify the number of puncta. Thresholds were set manually.

### Statistical analysis

Statistical comparisons were performed as such: Fisher’s exact test was used for categorical data between two groups (e.g. mistargeting, neuroblast glomerular innervation); Fisher- Freeman-Halton test was used for categorical data between three groups (e.g. mistargeting between controls, mutants, and rescues). Mann-Whitney U test was used to compare quantitative data between two groups (e.g. bouton diameter between controls and mutants); Kruskal-Wallis test and Dunn’s test post-hoc were used to compare quantitative data between three groups (e.g. bouton diameter between controls, mutants, and rescues). The numbers of independent replicates per experiment are indicated in the figures or legends. P-values were adjusted using the Benjamini-Hochberg procedure.

---

**Figure 1-figure supplement 1.**
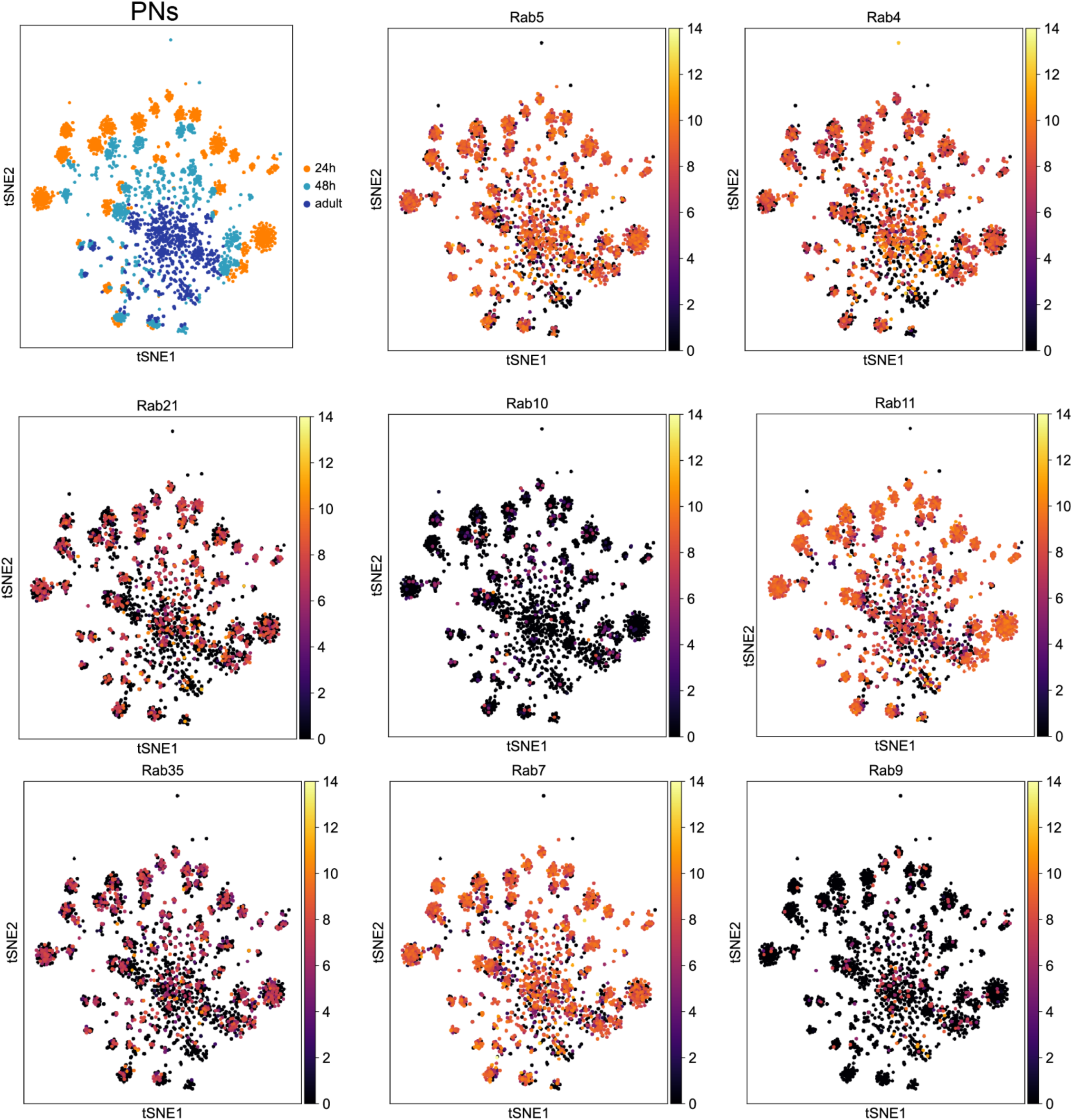
Expression of endosome-associated Rab GTPases in olfactory projection neurons. (Expression of Rab GTPases in developing PNs at 24h after puparium formation (APF) and 48h APF from single-cell RNA-seq (scRNA-seq) data. Expression is in log_2_(CPM +1), where CPM stands for transcript counts per million reads. scRNA-seq data are from Xie et al., 2021.

**Figure 1-figure supplement 2.**
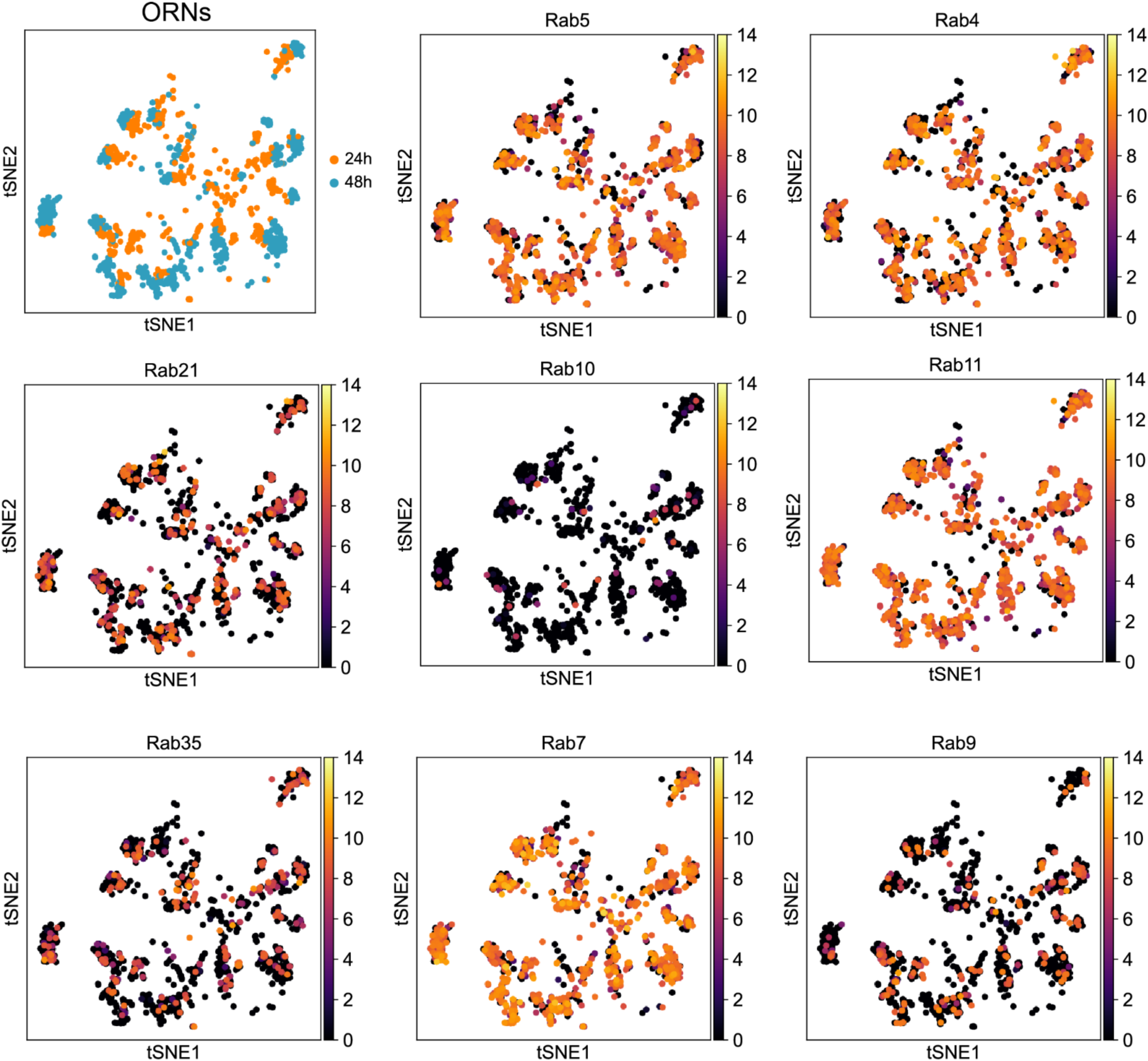
Expression of endosome-associated Rab GTPases in olfactory receptor neurons. Expression of Rab GTPases in developing ORNs at 24h after puparium formation (APF) and 48h APF from single-cell RNA-seq (scRNA-seq) data. Expression is in log_2_(CPM +1), where CPM stands for transcript counts per million reads. scRNA-seq data are from McLaughlin et al., 2021.

**Figure 1-figure supplement 3.**
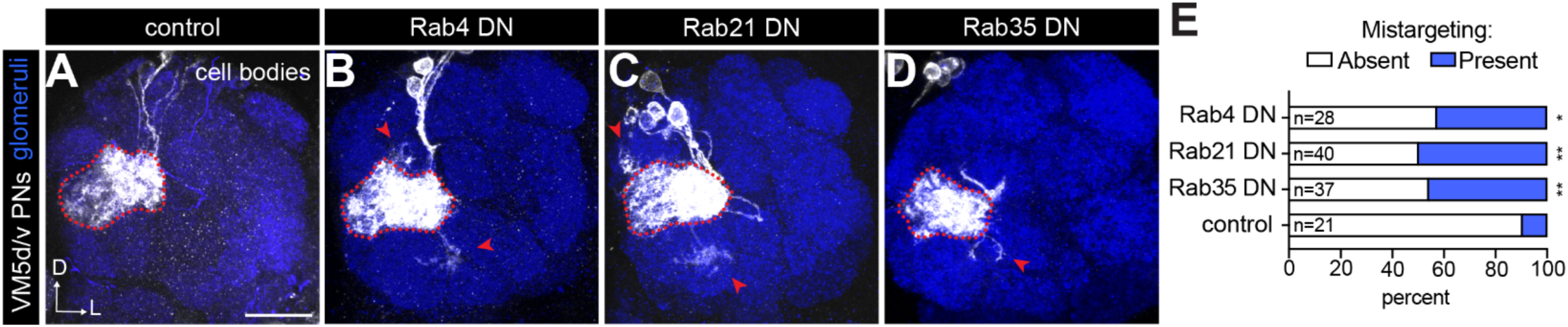
Extended analysis of endosome-associated Rab GTPases. (A–D) Representative images of indicated genotypes depicting phenotypes observed in dominant negative screen. Red dotted lines outline the VM5d/v glomeruli, and red arrows denote ectopic targeting. Scale bar, 20 μm. (F) Percent of antennal lobes with mistargeting in the Rab dominant negative screen. Fisher’s exact test compared to control.

**Figure 2-figure supplement 1.**
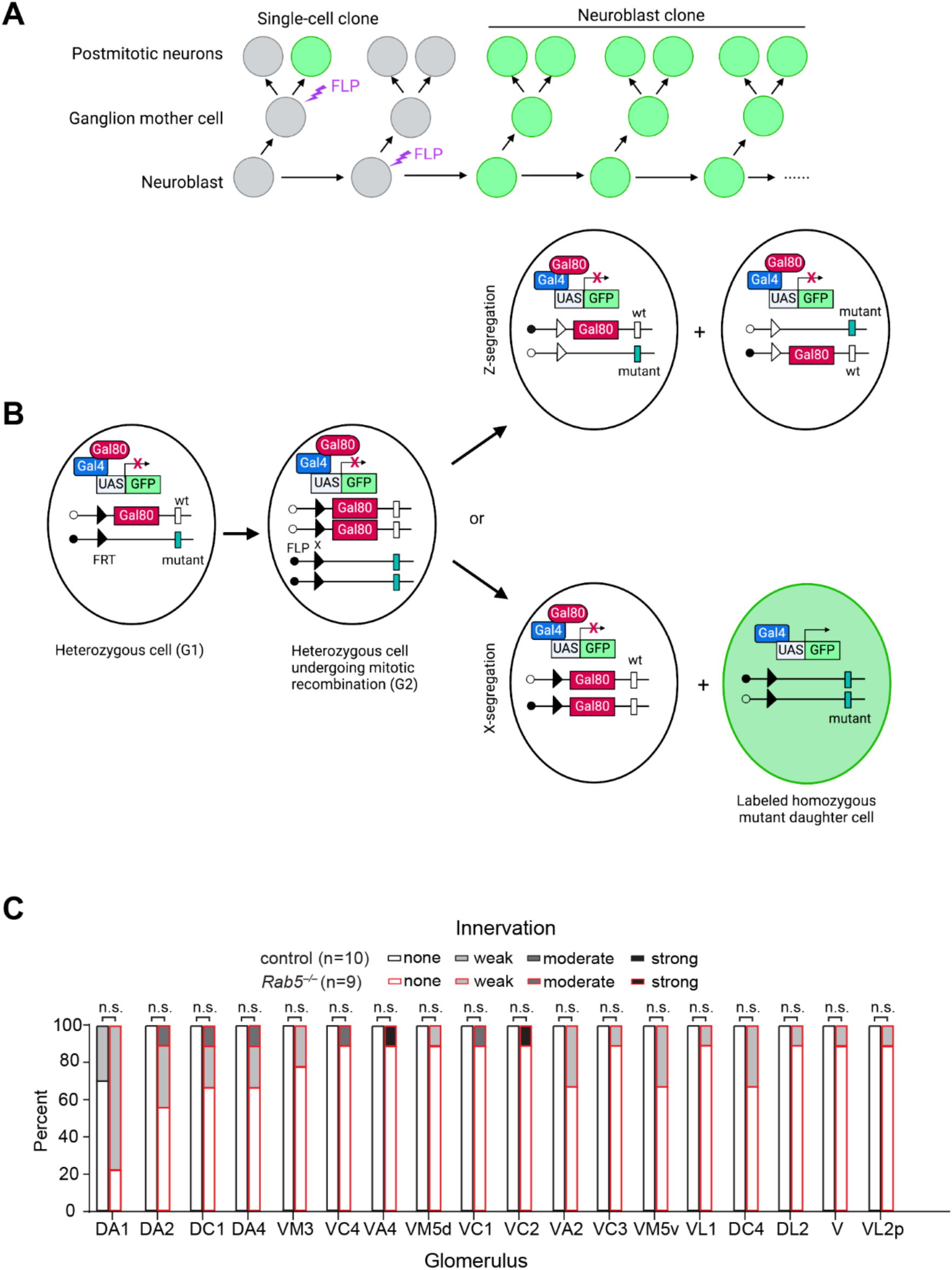
Schematic of MARCM-based mosaic analysis. (A) Schematic depicting heatshock flp induction of single-cell or neuroblast MARCM clones. MARCM can be used to generate GFP-labeled single-cell or neuroblast clones in PNs. All clones were induced by heat shock applied to newly hatched larvae (0–24h after larval hatching), so our analyses are primarily restricted to the adPNs and DL1-PN single-cell clones. (B) Schematic of MARCM analysis. A mutant (*Rab5*, *Rab7*, *Rab11*, or *Rab21*) allele is placed on a chromosome arm in *trans* to the chromosome arm with a *GAL80* transgene. Heterozygous cells express GAL80, which represses GAL4 activity and thus inhibits GFP expression in these cells. Following, FLP-mediated mitotic recombination and X-segregation (bottom row) one of the daughter cells becomes homozygous for the mutant allele and loses the *GAL80* transgene. Thus, homozygous mutant cells will be labeled with membrane-bound GFP (and can also express any UAS-based rescue transgene). (C) Quantification of adPN mistargeting in additional non-adPN glomeruli. Fisher’s exact test.

**Figure 6-figure supplement 1.**
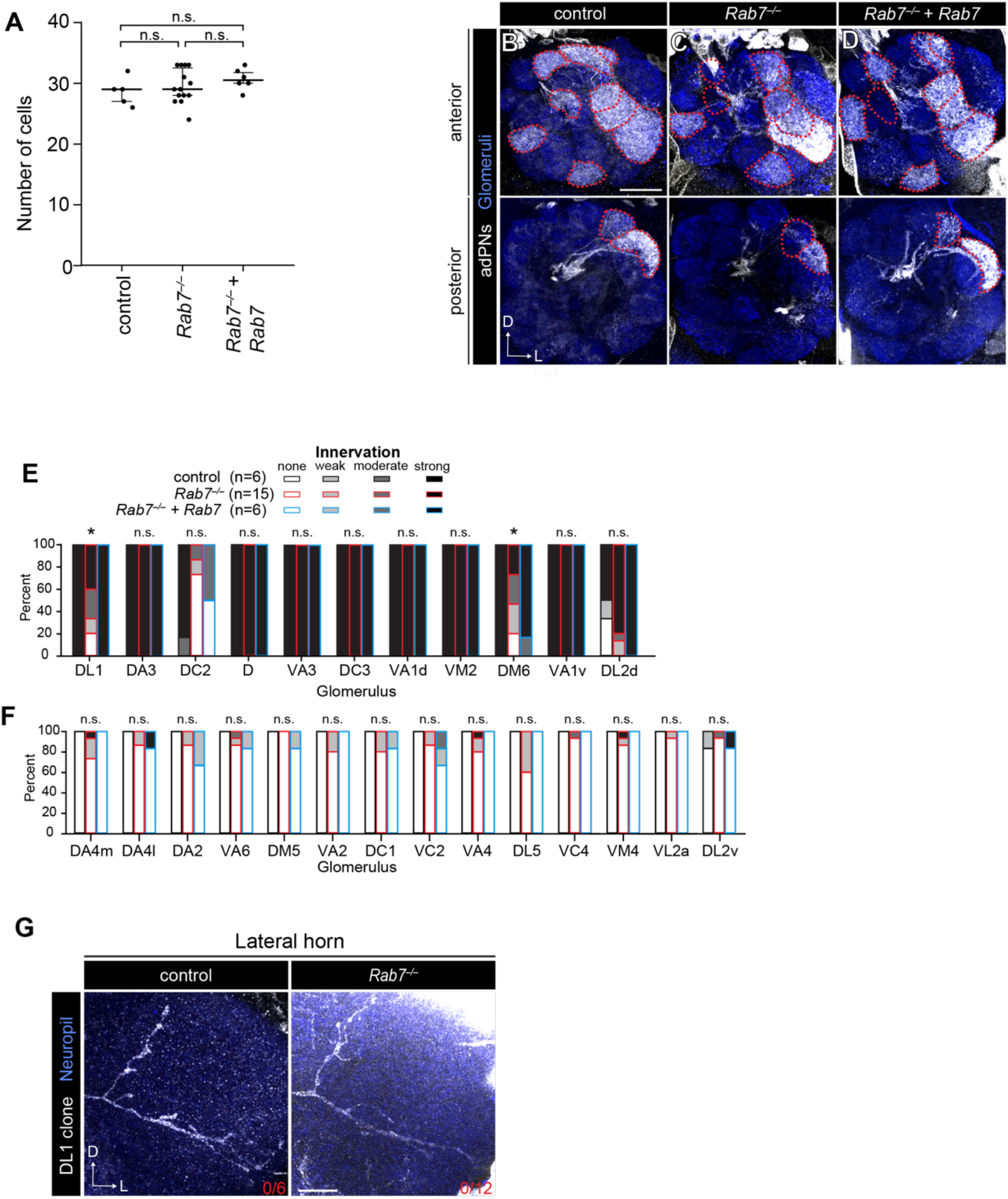
Extended analysis of *Rab7* loss-of-function MARCM phenotypes. (A) Quantification of number of cell bodies in adPN neuroblast clones of controls (n=5), *Rab7^−/−^* mutants (n=14), and *Rab7* rescues (n=6). Data are presented as medians (IQR). Kruskal-Wallis test followed by Dunn’s test post-hoc was used to assess statistical significance. (B–D) Representative images of adPN neuroblast clones of indicated genotypes. (E, F) Quantification of percent of antennal lobes with each category of dendrite innervation to adPN glomeruli (E) and non-adPN glomeruli (F) in indicated genotypes. Fisher-Freeman-Halton test was used to assess statistical significance. See Table S2 for complete statistical analysis. (G) Representative images of DL1-PN lateral horn axons in indicated genotypes. Scale bar, 20 µm (B), 10 µm (G).

**Figure 6-figure supplement 2.**
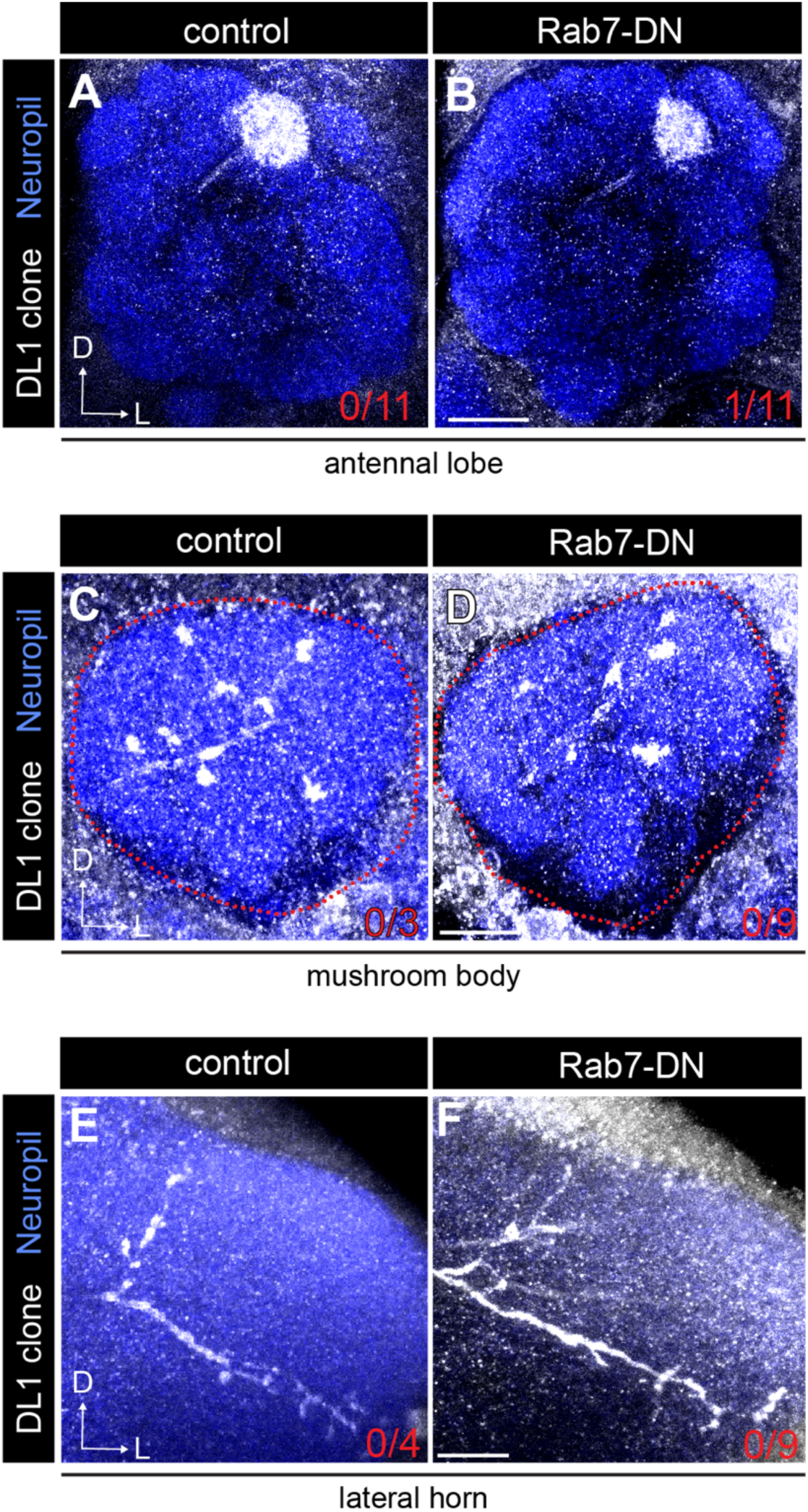
Rab7 is dispensable for DL1-PN development. (A, B) Representative images of DL1-PN dendrite targeting in indicated genotypes. Red numbers in the right corner of images denote DL1-PN dendrite mistargeting phenotypic penetrance. (C, D) Representative images of mushroom body axons in indicated genotypes. Red numbers in the right corner of images denote DL1-PN axon morphogenesis phenotypic penetrance. (E, F) Representative images of lateral axons in indicated genotypes. Red numbers in the right corner of images denote DL1-PN axon morphogenesis phenotypic penetrance. Scale bar, 20 µm (C); 10 µm (D, I)

**Figure 7-figure supplement 1.**
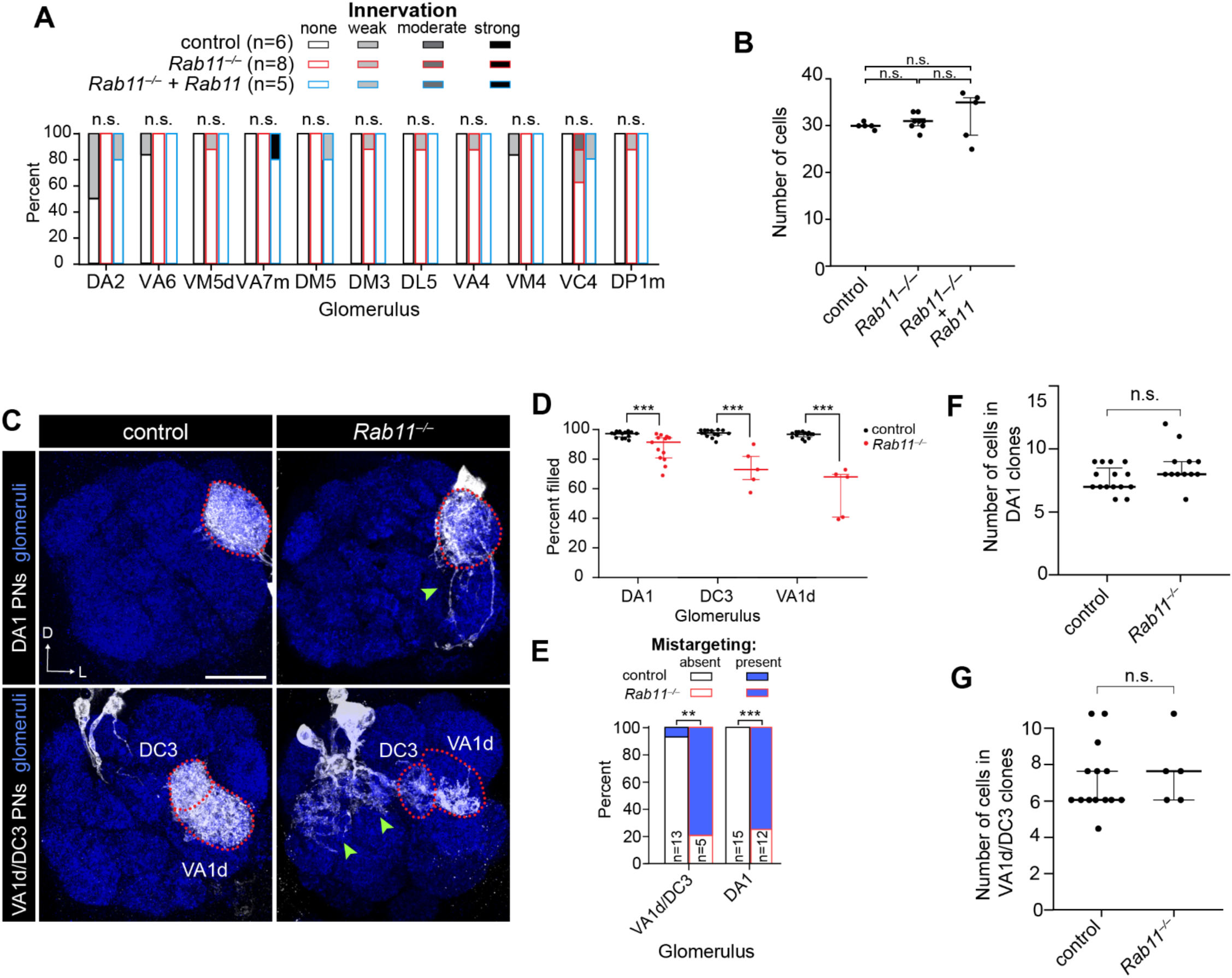
Additional analysis of *Rab11* dendrite targeting phenotypes. (A) Quantification of ectopic dendrite targeting phenotypes across all analyzed non-adPN glomeruli, excluding those present in Figure 5E. Note the non-adPN glomeruli present in Figure 5E are significantly different. Fisher-Freeman-Halton test was used to assess statistical significance. See Table S3 for complete statistical analyses. (B) Quantification of number of cell bodies in adPN neuroblast clones of controls (n=5), *Rab11*^−/−^ mutants (n=8), and *Rab11* rescues (n=5). Data are presented as medians (IQR). Kruskal-Wallis test followed by Dunn’s test post-hoc test was used to assess statistical significance. (C) Representative images of mistargeting observed in indicated genotypes. Red dotted outline denotes DA1 (top row) or DC3/VA1d (bottom row) glomeruli. Green arrows denote mistargeting. (D) Quantification of the percent of the indicated glomerulus filled with PN dendrites in DA1-PN controls (n=15) and *Rab11*^−/−^ mutants (n=12) and VA1d/DC3-PN controls (n=14) and *Rab11*^−/−^ mutants (n=5). Data are presented as medians (IQR). Mann-Whitney U test was used to assess statistical significance. (E) Quantification of the proportion of antennal lobes with mistargeting. Fisher’s exact test was used to assess statistical significance. (F) Quantification of the number of cell bodies in neuroblast clones containing labeled DA1-PNs visualized by the MZ19-GAL4 driver for controls (n=15) and *Rab11*^−/−^ mutants (n=12). Data are presented as medians (IQR). Mann-Whitney U test was used to determine assess significance. (G) Quantification of the number of cell bodies in neuroblast clones containing labeled VA1d/DC3-PNs visualized by the MZ19-GAL4 driver controls (n=14) and *Rab11*^−/−^ mutants (n=5). Data are presented as medians (IQR). Mann-Whitney U test was used to assess statistical significance. Scale bar, 20 µm (C).

**Table S1.**
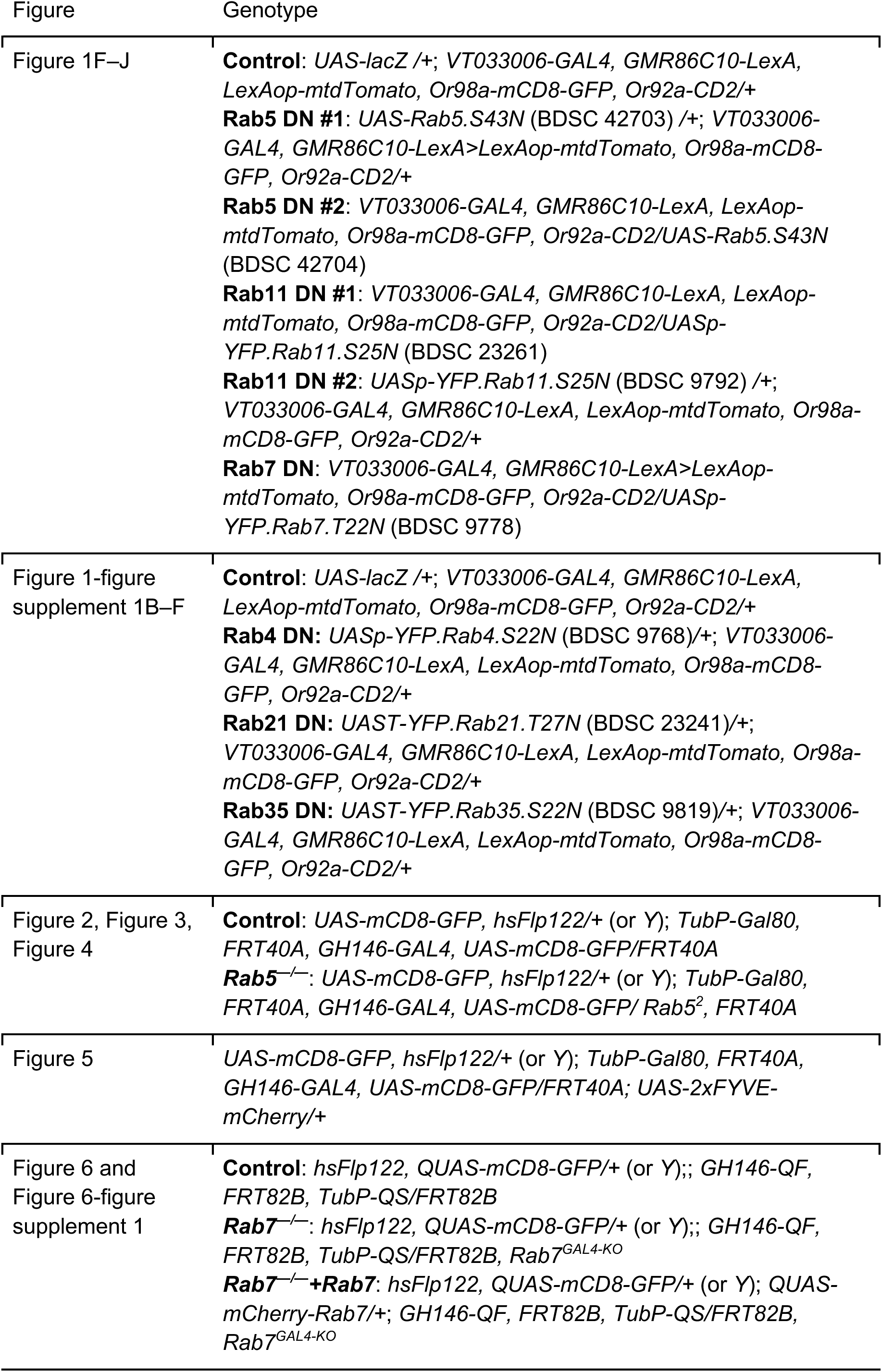
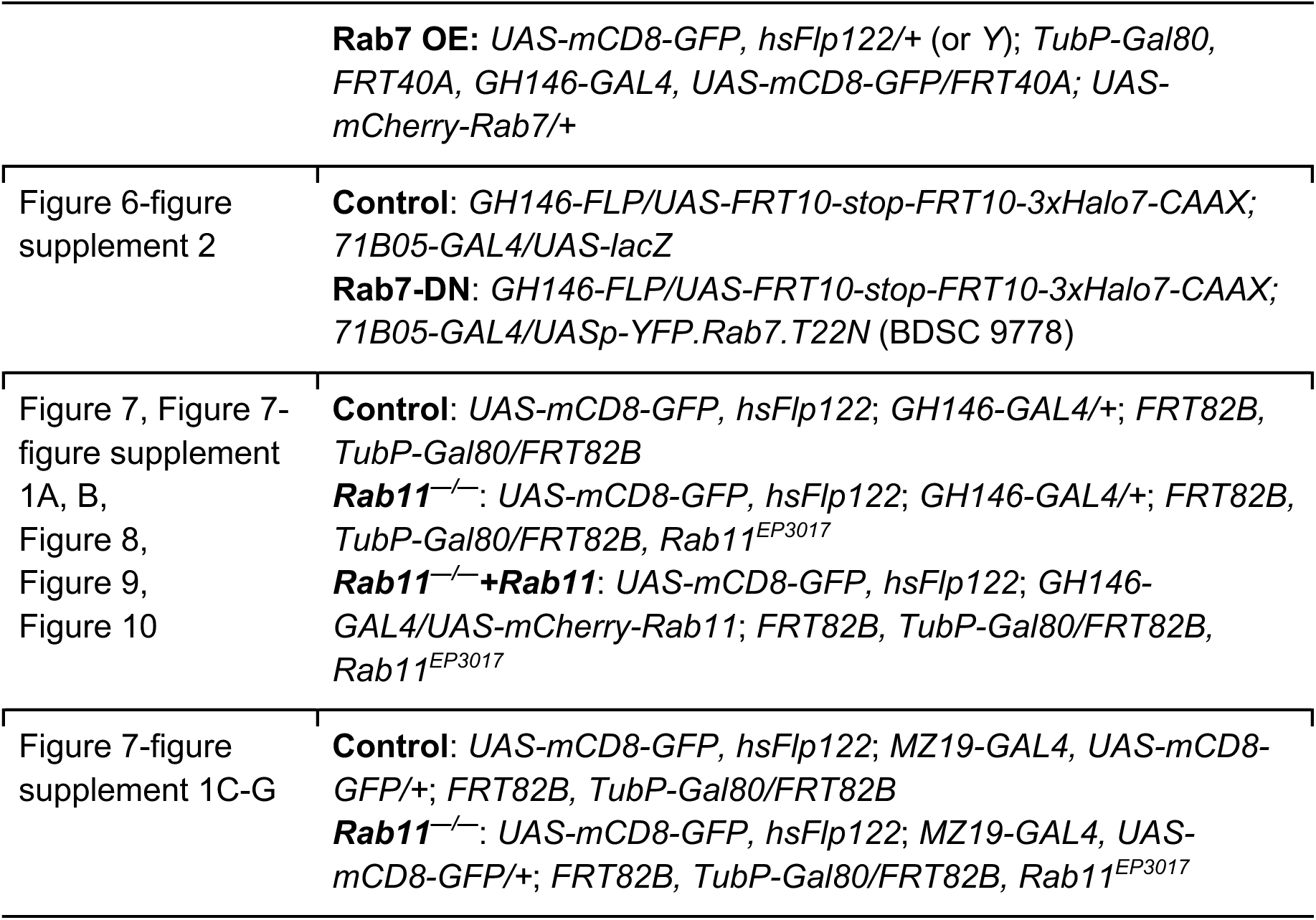
Complete list of genotypes used in this study.

| Figure | Genotype |
| --- | --- |
| Figure 1F–J | **Control:** *UAS-lacZ* /+; *VT033006-GAL4*, *GMR86C10-LexA*, *LexAop-mtdTomato*, *Or98a-mCD8-GFP*, *Or92a-CD2*/+<br>**Rab5 DN #1:** *UAS-Rab5.S43N* (BDSC 42703) /+; *VT033006-GAL4*, *GMR86C10-LexA*>*LexAop-mtdTomato*, *Or98a-mCD8-GFP*, *Or92a-CD2*/+<br>**Rab5 DN #2:** *VT033006-GAL4*, *GMR86C10-LexA*, *LexAop-mtdTomato*, *Or98a-mCD8-GFP*, *Or92a-CD2/UAS-Rab5.S43N* (BDSC 42704)<br>**Rab11 DN #1:** *VT033006-GAL4*, *GMR86C10-LexA*, *LexAop-mtdTomato*, *Or98a-mCD8-GFP*, *Or92a-CD2/UASp-YFP.Rab11.S25N* (BDSC 23261)<br>**Rab11 DN #2:** *UASp-YFP.Rab11.S25N* (BDSC 9792) /+; *VT033006-GAL4*, *GMR86C10-LexA*, *LexAop-mtdTomato*, *Or98a-mCD8-GFP*, *Or92a-CD2*/+<br>**Rab7 DN:** *VT033006-GAL4*, *GMR86C10-LexA*>*LexAop-mtdTomato*, *Or98a-mCD8-GFP*, *Or92a-CD2/UASp-YFP.Rab7.T22N* (BDSC 9778) |
| Figure 1-figure supplement 1B–F | **Control:** *UAS-lacZ* /+; *VT033006-GAL4*, *GMR86C10-LexA*, *LexAop-mtdTomato*, *Or98a-mCD8-GFP*, *Or92a-CD2*/+<br>**Rab4 DN:** *UASp-YFP.Rab4.S22N* (BDSC 9768)/+; *VT033006-GAL4*, *GMR86C10-LexA*, *LexAop-mtdTomato*, *Or98a-mCD8-GFP*, *Or92a-CD2*/+<br>**Rab21 DN:** *UAST-YFP.Rab21.T27N* (BDSC 23241)/+; *VT033006-GAL4*, *GMR86C10-LexA*, *LexAop-mtdTomato*, *Or98a-mCD8-GFP*, *Or92a-CD2*/+<br>**Rab35 DN:** *UAST-YFP.Rab35.S22N* (BDSC 9819)/+; *VT033006-GAL4*, *GMR86C10-LexA*, *LexAop-mtdTomato*, *Or98a-mCD8-GFP*, *Or92a-CD2*/+ |
| Figure 2, Figure 3, Figure 4 | **Control:** *UAS-mCD8-GFP*, *hsFlp122*/+ (or *Y*); *TubP-Gal80*, *FRT40A*, *GH146-GAL4*, *UAS-mCD8-GFP/FRT40A*<br>**Rab5<sup>-/-</sup>:** *UAS-mCD8-GFP*, *hsFlp122*/+ (or *Y*); *TubP-Gal80*, *FRT40A*, *GH146-GAL4*, *UAS-mCD8-GFP/Rab5<sup>2</sup>*, *FRT40A* |
| Figure 5 | *UAS-mCD8-GFP*, *hsFlp122*/+ (or *Y*); *TubP-Gal80*, *FRT40A*, *GH146-GAL4*, *UAS-mCD8-GFP/FRT40A*; *UAS-2xFYVE-mCherry*/+ |
| Figure 6 and Figure 6-figure supplement 1 | **Control:** *hsFlp122*, *QUAS-mCD8-GFP*/+ (or *Y*); *GH146-QF*, *FRT82B*, *TubP-QS/FRT82B*<br>**Rab7<sup>-/-</sup>:** *hsFlp122*, *QUAS-mCD8-GFP*/+ (or *Y*); *GH146-QF*, *FRT82B*, *TubP-QS/FRT82B*, *Rab7<sup>GAL4-KO</sup>*<br>**Rab7<sup>-/-</sup>+Rab7:** *hsFlp122*, *QUAS-mCD8-GFP*/+ (or *Y*); *QUAS-mCherry-Rab7*/+; *GH146-QF*, *FRT82B*, *TubP-QS/FRT82B*, *Rab7<sup>GAL4-KO</sup>* |
|  | **Rab7 OE:** *UAS-mCD8-GFP, hsFlp122/+ (or Y); TubP-Gal80, FRT40A, GH146-GAL4, UAS-mCD8-GFP/FRT40A; UAS-mCherry-Rab7/+* |
| Figure 6-figure supplement 2 | **Control:** *GH146-FLP/UAS-FRT10-stop-FRT10-3xHalo7-CAAX; 71B05-GAL4/UAS-lacZ*<br>**Rab7-DN:** *GH146-FLP/UAS-FRT10-stop-FRT10-3xHalo7-CAAX; 71B05-GAL4/UASp-YFP.Rab7.T22N (BDSC 9778)* |
| Figure 7, Figure 7-figure supplement 1A, B, Figure 8, Figure 9, Figure 10 | **Control:** *UAS-mCD8-GFP, hsFlp122; GH146-GAL4/+; FRT82B, TubP-Gal80/FRT82B*<br>**Rab11<sup>-/-</sup>:** *UAS-mCD8-GFP, hsFlp122; GH146-GAL4/+; FRT82B, TubP-Gal80/FRT82B, Rab11<sup>EP3017</sup>*<br>**Rab11<sup>-/-</sup>+Rab11:** *UAS-mCD8-GFP, hsFlp122; GH146-GAL4/UAS-mCherry-Rab11; FRT82B, TubP-Gal80/FRT82B, Rab11<sup>EP3017</sup>* |
| Figure 7-figure supplement 1C-G | **Control:** *UAS-mCD8-GFP, hsFlp122; MZ19-GAL4, UAS-mCD8-GFP/+; FRT82B, TubP-Gal80/FRT82B*<br>**Rab11<sup>-/-</sup>:** *UAS-mCD8-GFP, hsFlp122; MZ19-GAL4, UAS-mCD8-GFP/+; FRT82B, TubP-Gal80/FRT82B, Rab11<sup>EP3017</sup>* |

**Table S2.**
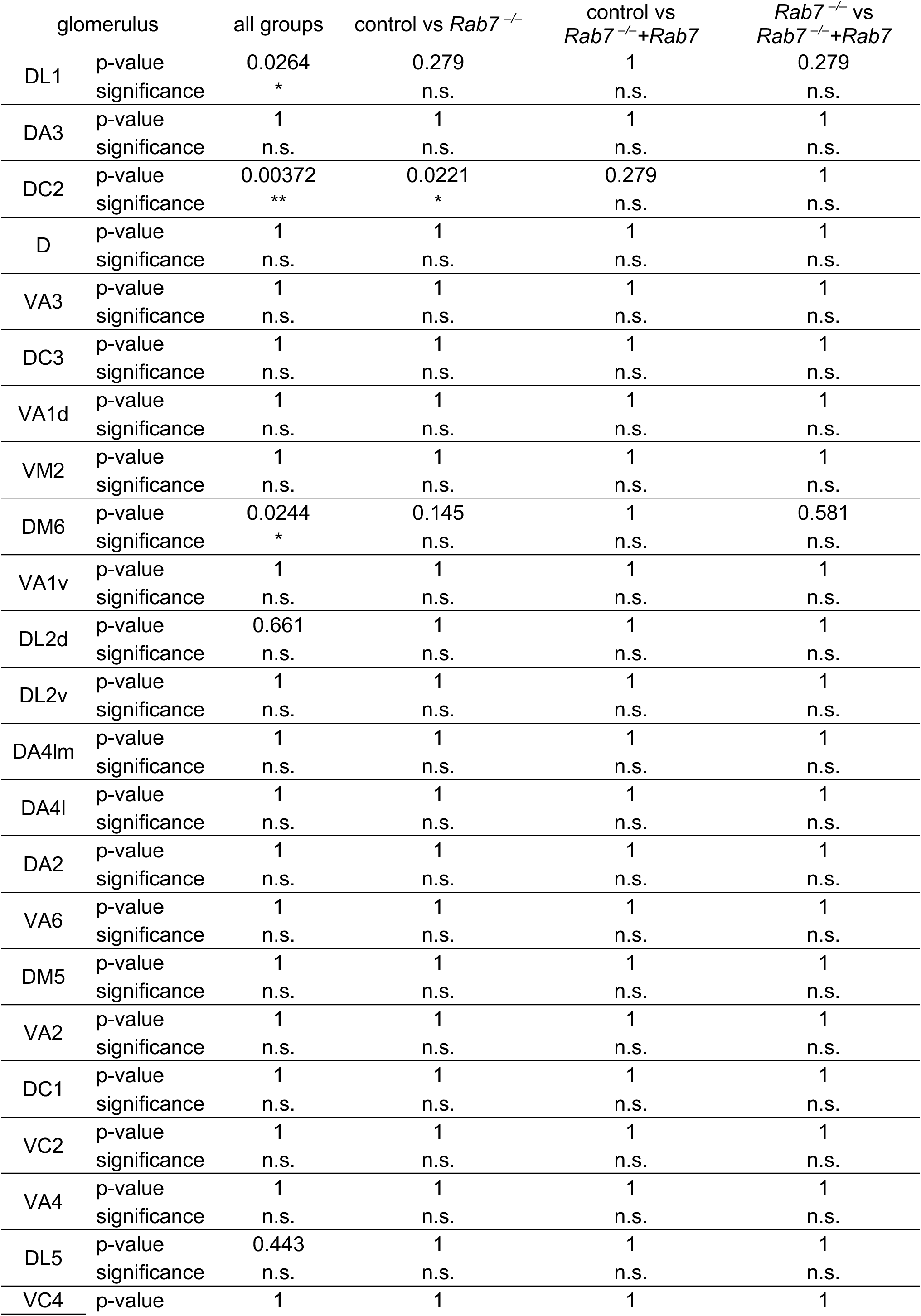
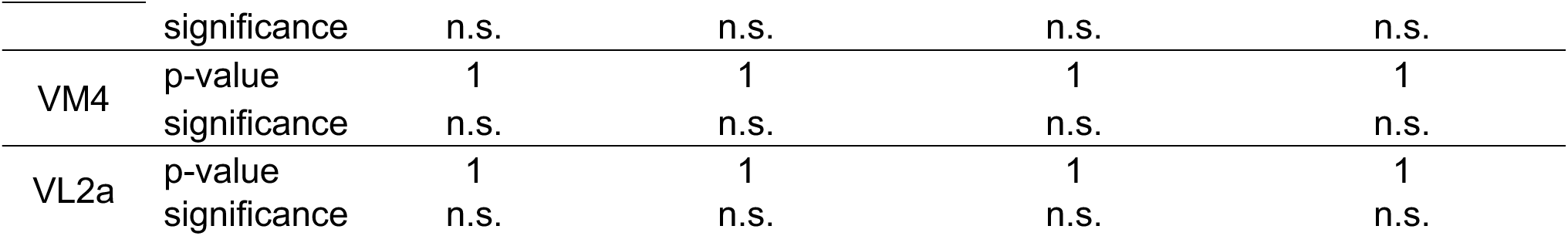
Complete statistical analysis for *Rab7* mutant adPN neuroblast clones. Statistical significance was assessed using the Fisher-Freeman-Halton test for overall group comparisons, followed by pairwise Fisher’s exact tests with Benjamini-Hochberg correction for multiple comparisons.

**Table S3.**
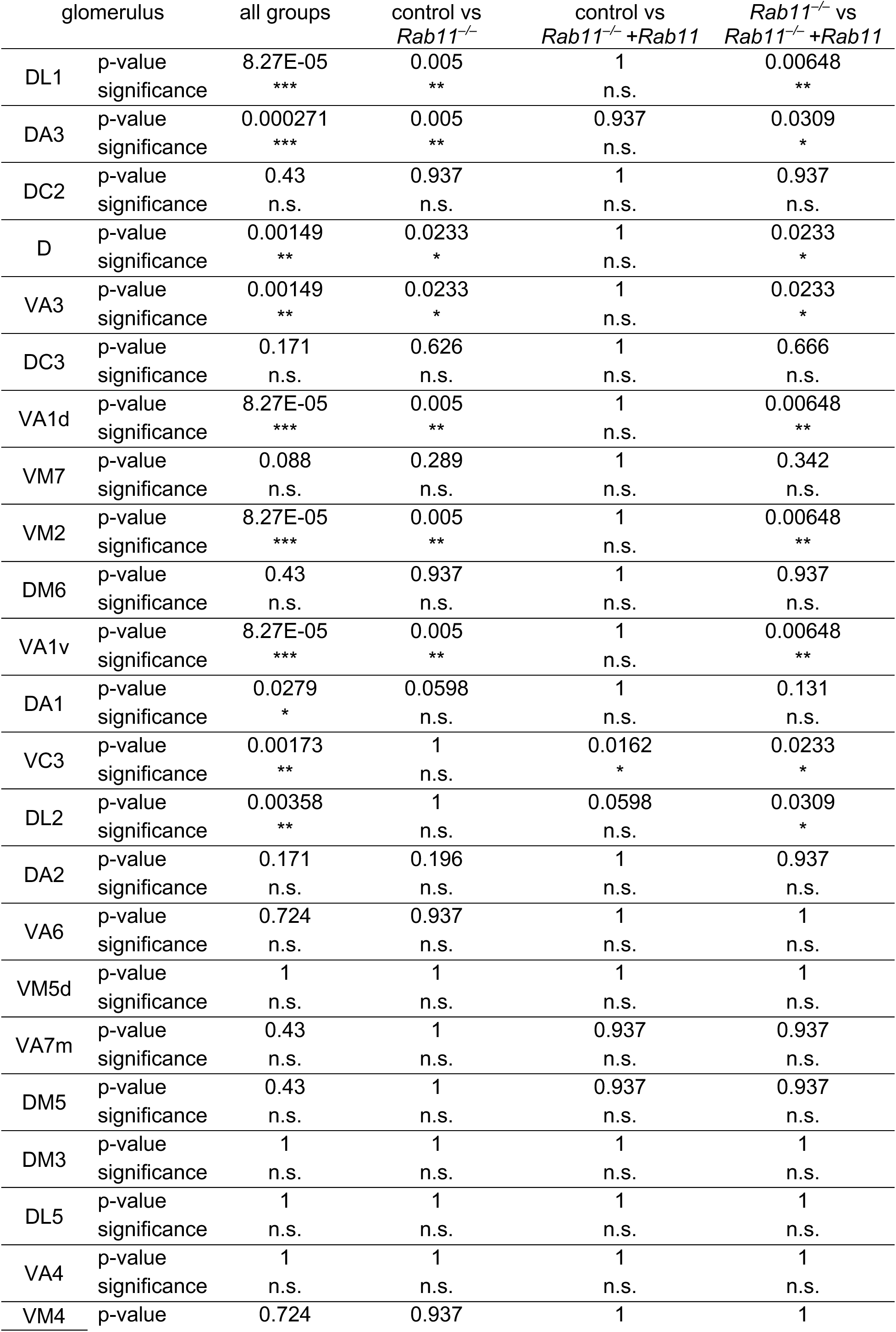
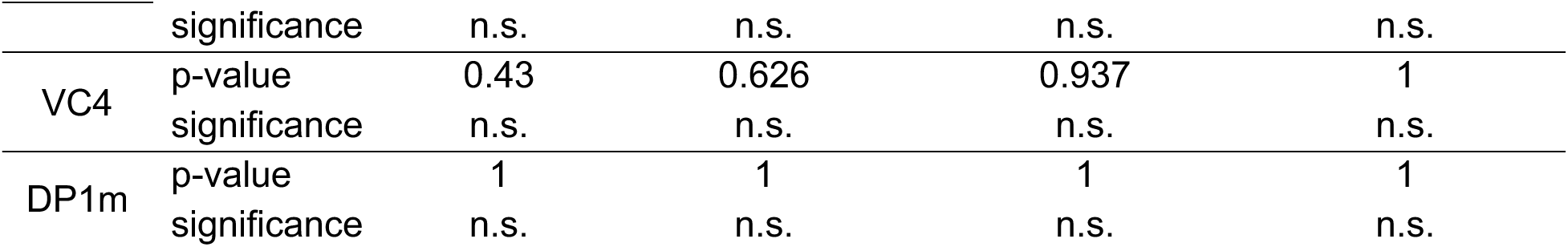
Complete statistical analysis for *Rab11* mutant adPN neuroblast clones. Statistical significance was assessed using the Fisher-Freeman-Halton test for overall group comparisons, followed by pairwise Fisher’s exact tests with Benjamini-Hochberg correction for multiple comparisons.

## Notes

### Competing Interest Statement

The authors have declared no competing interest.

### Summary of Updates

This manuscript has undergone peer-review has has been revised to incorporate reviewer comments.

